# Accurate model of liquid-liquid phase behaviour of intrinsically-disordered proteins from optimization of single-chain properties

**DOI:** 10.1101/2021.06.23.449550

**Authors:** Giulio Tesei, Thea K. Schulze, Ramon Crehuet, Kresten Lindorff-Larsen

## Abstract

Many intrinsically disordered proteins (IDPs) may undergo liquidliquid phase separation (LLPS) and participate in the formation of membraneless organelles in the cell, thereby contributing to the regulation and compartmentalisation of intracellular biochemical reactions. The phase behaviour of IDPs is sequence-dependent, and its investigation through molecular simulations requires protein models that combine computational efficiency with an accurate description of intra- and intermolecular interactions. We developed a general coarse-grained model of IDPs, with residue-level detail, based on an extensive set of experimental data on single-chain properties. Ensemble-averaged experimental observables are predicted from molecular simulations, and a data-driven parameter-learning procedure is used to identify the residue-specific model parameters that minimize the discrepancy between predictions and experiments. The model accurately reproduces the experimentally observed conformational propensities of a set of IDPs. Through two-body as well as large-scale molecular simulations, we show that the optimization of the intramolecular interactions results in improved predictions of protein self-association and LLPS.

---

Many intrinsically disordered proteins (IDPs) and proteins with disordered regions can condense into liquid-like droplets, viz. a biomolecule-rich phase coexisting with a more dilute solution (1–5). This de-mixing process is known as liquid-liquid phase separation (LLPS) and is one of the ways cells compartmentalise proteins, often together with nucleic acids (6). While LLPS plays crucial biological roles in the cell, its dysregulation leads to maturation of biomolecular condensates into hydrogel-like assemblies, promoting the formation of neurotoxic oligomers and amyloid fibrils (5, 7). A quantitative model for the ‘molecular grammar’ of LLPS, including the influence of disease-associated mutations and post-translational modifications (PTMs) on the propensity to phase separate, is key to understand these processes. The sequences of IDPs and intrinsically disordered regions that easily undergo LLPS are often characterized by stretches enriched in small polar residues (spacers) interspersed by e.g. aromatic or arginine residues (stickers), which are instrumental for the formation of reversible physical cross-links via *π*-*π*, cation-*π* and sp^2^-*π* interactions (8–12). Y and R residues were shown to be necessary for the LLPS of a number of proteins including FUS, hnRNPA1, LAF-1 and Ddx4 (8,10,11,13–17). While the propensity to undergo LLPS increases with the number of Y residues in the sequence, recent studies have revealed that the role of R residues is context dependent (16) and strongly affected by salt concentration (17), reflecting the unusual characteristics of the R side chain (18, 19).

Here, we present the development of a coarse-grained (CG) model capable of predicting the phase behaviour of IDPs based on amino acid sequence. CG models enable the combination of a sequence-dependent description with the computational efficiency necessary to explore the long time and large length scales involved in phase transitions (11,20,21). Although CG molecular simulations have been employed to explain the sequence dependence of the LLPS of a number of IDPs (11,15,17,20–22) as well as the effect of phosphorylation on LLPS propensities (23, 24), such models have proven difficult to use to predict the phase behaviour of very diverse sequences (25). Building on recent developments, including experimental phase diagrams of a number of IDPs (3,4,15,16), we trained and tested a robust sequence-dependent model of the LLPS of IDPs. In particular, due to the similarity between intramolecular interactions within IDPs and intermolecular interactions between IDPs (12, 26), we rationalized that by optimizing a model to capture structural preferences for a broad set of monomeric IDPs, we could obtain a good model for interactions between IDPs.

The starting point for our analyses is the hydrophobicity scale (HPS) model (21) (with minor modification; see *SI Appendix*) wherein, besides steric repulsion and salt-screened charge-charge interactions, residue-residue interactions are determined by hydropathy parameters (λ) which were derived from the atomic partial charges of a classical all-atom force field (27). Recently, the development of the HPS-Urry model (28) presented substantial improvements in accuracy over the original HPS model. These were achieved using a hydrophobicity scale derived from transition temperatures of elastin-like peptides (29), and further shifting the λ parameters by −0.08 to improve agreement with experimentally measured radii of gyration.

To address the current limitations, we improve upon these models by optimizing the λ parameters through a Bayesian parameter-learning procedure (30–33), leveraging as prior knowledge the probability distribution of the λ parameters evaluated from analysing 87 hydrophobicity scales. The training set comprises SAXS and paramagnetic relaxation enhancement (PRE) NMR data of 45 IDPs which we selected from the literature. First, we run Langevin dynamics simulations of single IDPs and estimate the experimental observables using state-of-the-art methods (34). Second, we employ a Bayesian regularization approach to prevent over-fitting the training data and select three models which are equally accurate with respect to single-chain conformational properties. Third, through two-chain simulations, we validate the models by comaparing predicted and experimental intermolecular PRE NMR data for the low complexity domain (LCD) of the heterogeneous nuclear ribonucleoprotein (hnRNP) A2 (A2 LCD) (22) and the LCD of the RNA-binding protein fused in sarcoma (FUS LCD) (23). Fourth, we perform coexistence simulations to test the models against the phase behaviour of A2 LCD (22, 24), FUS LCD (35, 36), variants of hnRNP A1 LCD (A1 LCD) (15, 16), the N-terminal region of the germ-granule protein Ddx4 (Ddx4 LCD) (8,10,13) and the N-terminal, R-/G-rich domain of the P granule protein LAF-1 (LAF-1 RGG domain). We use the final model to provide insight into the interactions between IDPs within condensates and to help elucidate the role of different amino acids to the driving force for LLPS.

## Results and Discussion

### Analysis of Hydrophobicity Scales

The λ values of the original HPS model are based on a hydrophobicity scale derived by Kapcha and Rossky from the atomic partial charges of the OPLS all-atom force field (27). Dozens of amino acid hydrophobicity scales have been derived from experimental as well as bioinformatics approaches such as the partitioning of amino acids between water and organic solvent, the partitioning of peptides to the lipid membrane interface and the accessible surface area of residues in folded proteins (38, 39). To carry out the Bayesian optimisation of the amino-acid specific λ values, we sought to estimate the prior probability distribution of the hydropathy parameters from the analysis of 98 hydrophobicity scales collected by Simm et al. (39). Each scale was min-max normalized and, after ranking in the ascending order of the HPS scale, we discarded all the scales yielding a linear fit with negative slope. This procedure allowed us to identify scales which were present in the set both in their original form and as the additive inverse of the hydropathy values (reversed scales). For most scales, the selection criterion resulted in discarding the reversed form. However, for scales where the most negative values of the hydropathy parameter correspond to the most hydrophobic amino acids—such as the scales by Bull and Breese (40), Guy (41), Bishop et al. (42) and Welling et al. (43)—we retained only the reversed form. The 87 scales that remained after this filtering were used to calculate the average scale (AVG) and the probability distribution of the λ values for the 20 amino acids, *P*(λ), which is normalized so that 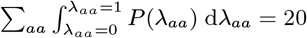 (Fig. 1*A*). For the optimization described below we use the AVG scale as starting point, as well as an indication of the typical accuracy obtained from the prior knowledge encoded in *P*(λ).

**Fig. 1.**
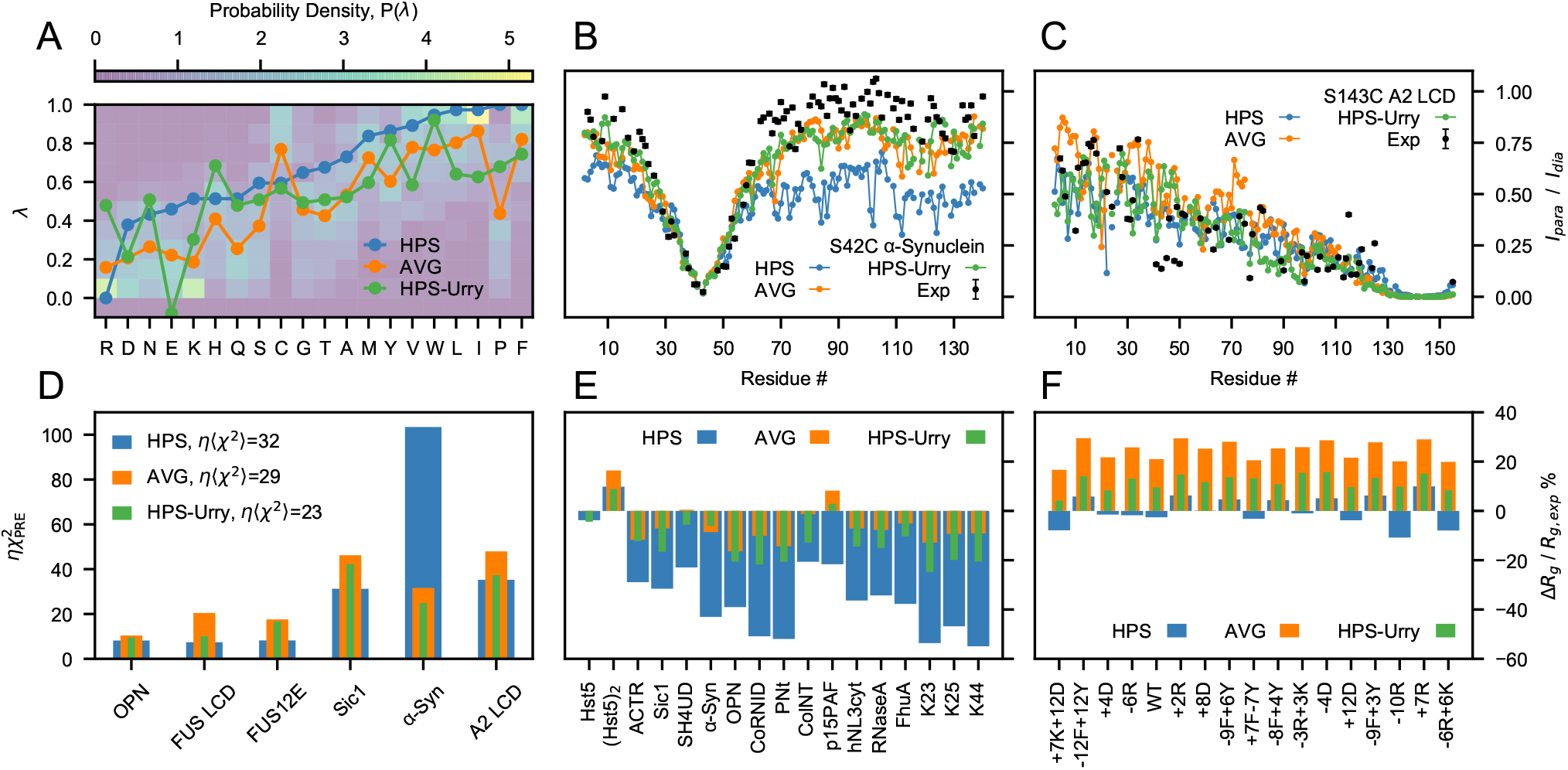
Assessing the HPS, AVG and HPS-Urry models using experimental data reporting on single-chain conformational properties. (*A*) Probability distributions of the λ parameters calculated from 87 min-max normalized hydrophobicity scales. Lines are the λ parameters of the HPS model (blue), the average over the hydrophobicity scales (orange) and the HPS-Urry model (green) (28). Intramolecular PRE intensity ratios for the S43C mutant of *α*-Synuclein (*B*) and the S243C mutant of A2 LCD (*C*) from simulations and experiments (22, 37) (black). (*D*) *χ*^2^ values quantifying the discrepancy between simulated and experimental intramolecular PRE data, scaled by the hyperparameter *η* = 0.1 (Materials and Methods). *(E* and *F*) Relative difference between simulated and experimental radii of gyration for proteins that do not readily undergo phase separation alone (*E*) and for variants of A1 LCD (*F*), with negative values corresponding to the simulated ensembles being more compact than in experiments.

We assessed the HPS, HPS-Urry and AVG parameter sets by running simulations of 45 IDPs ranging in length between 24 and 334 residues and compared the results against experiments. Specifically, we compared the simulations with the radii of gyration, *R_g_*, of 42 IDPs (Tab. S1) and intramolecular PRE data of six IDPs (Tab. S2) (16,22,23,37,44–57). Compared to the AVG scale, the HPS model overestimates the compaction of *α*-Synuclein whereas it closely reproduces the PRE data for A2 LCD (Fig. 1*B* and *C*). In general, the HPS model accurately predicts the conformational properties of sequences with high LLPS propensity, e.g. FUS LCD, A2 LCD and A1 LCD (Fig. 1*D* and *F*), while the AVG scale is considerably more accurate at reproducing the *R_g_* of proteins that do not readily undergo phase separation alone (Fig. 1*E*). The recently proposed HPS-Urry model (28) is the most accurate at predicting the intramolecular PRE data while it shows intermediate accuracy for the *R_g_* values of both proteins that do not readily undergo phase separation alone and A1 LCD variants. The HPS-Urry model in particular differs significantly from the HPS and AVG models for the λ parameters for R and E as well as the reversal of the order of hydrophobicity of Y and F (Fig. 1*A*).

### Optimization of Amino-Acid Specific Hydrophobicity Values

To obtain a model that more accurately predicts the conformational properties of IDPs of diverse sequences and LLPS propensities, we trained the λ values on a large set of experimental *R_g_* and PRE data using a Bayesian parameter-learning procedure (30) shown schematically in Fig. 2 (Materials and Methods). We initially performed an optimization run starting from the AVG λ values and setting the hyperparameters to *θ* = *η* = 0.1 (Fig. S1*A*). We collected the optimized sets of λ values which yielded 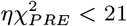 and 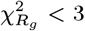 (circles in Fig. 3*A*). The optimization was repeated starting from all λ = 0.5 to assess that the parameter space sampled by our method is independent of the initial conditions (Fig. S2*A* and S1*D*). Thus, while we used the AVG model as starting point, our final parameters only depend on *P*(λ) via its use as the prior in the Bayesian optimization.

**Fig. 2.**
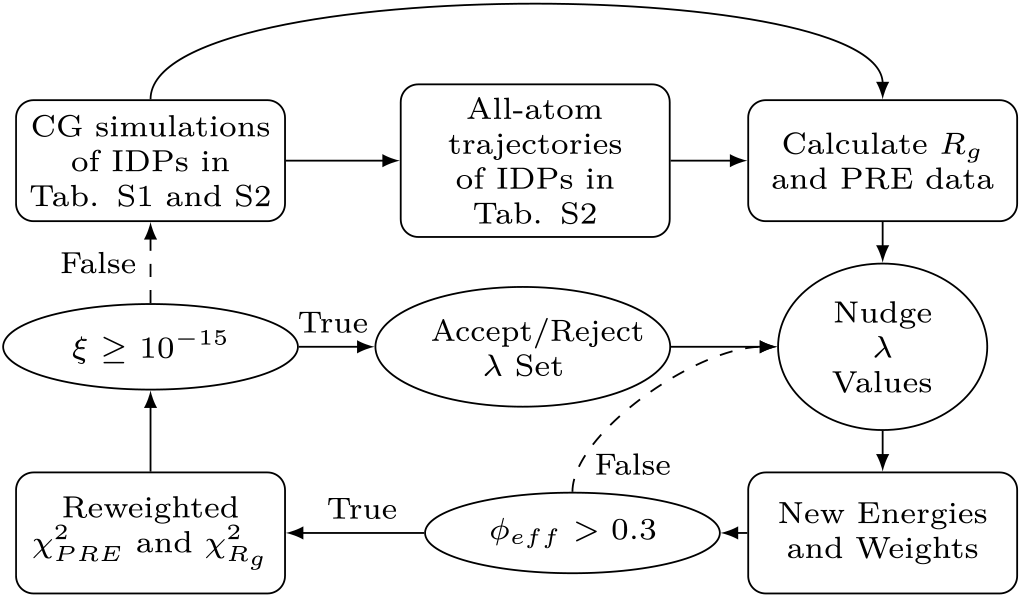
Flowchart illustrating the Bayesian parameter-learning procedure (Materials and Methods).

**Fig. 3.**
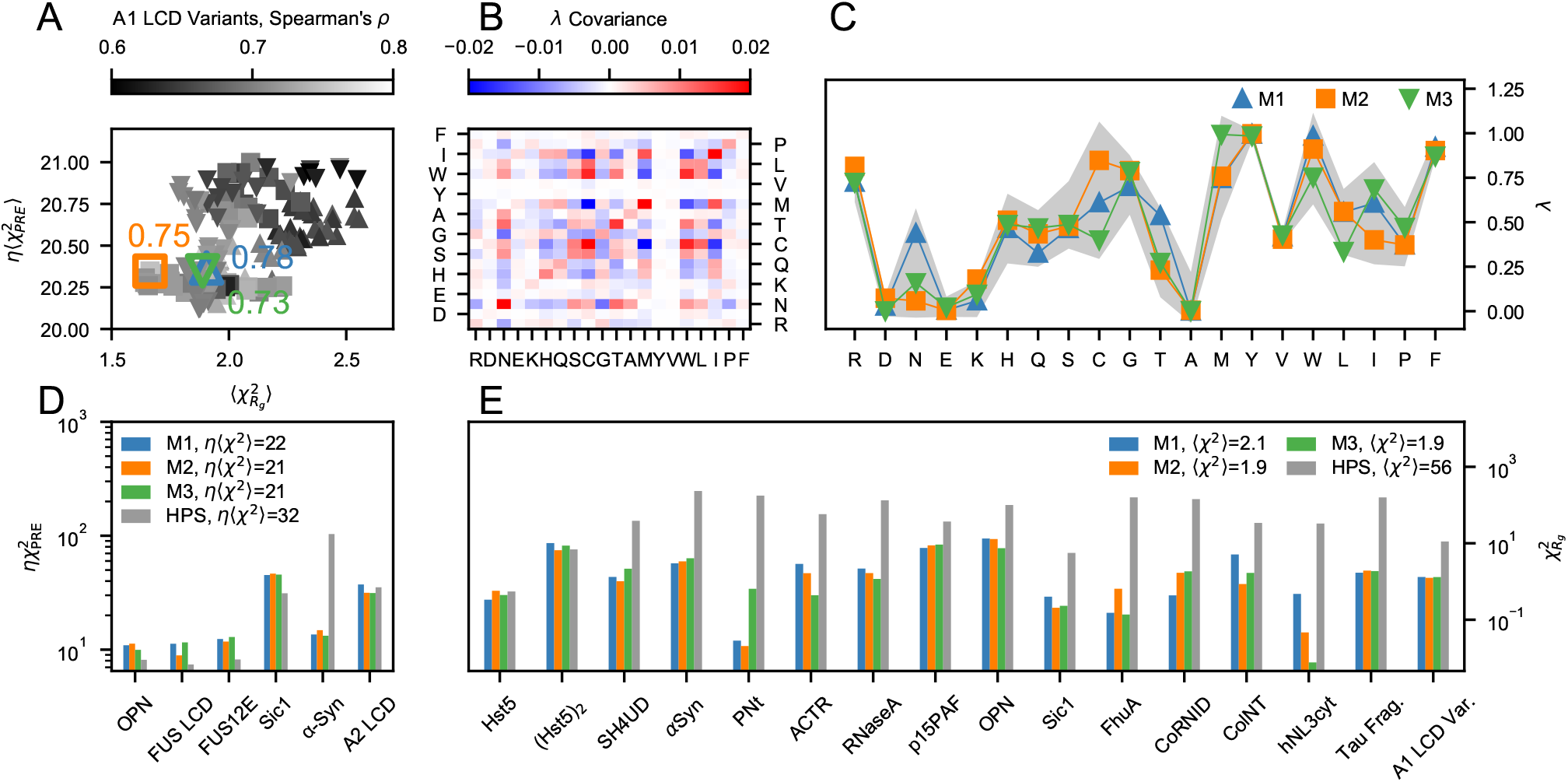
Selection and performance of the M1–3 models with respect to the training data. (*A*) Overview of the optimal λ sets with 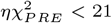 and 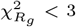 collected through the parameter learning procedures started from λ_0_ =AVG (up triangles), M1 (squares) and M2 (down triangles). The gray gradient shows the Spearman’s correlation coefficient between experimental and simulated *R_g_* values for the A1 LCD variants in the training set. Colored open symbols indicate the M1 (blue up triangle), M2 (orange square) and M3 (green down triangle) scales whereas the adjacent values are the respective Spearman’s correlation coefficients. (*B*) Covariance matrix of the λ sets with 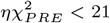 and 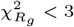. (*C*) M1 (blue), M2 (orange) and M3 (green) scales. Solid lines are guides for the eye whereas the gray shaded area shows the mean ±2SD of the λ sets with 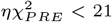 and 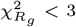. (*D*–*E*) Comparison between (*D*) 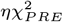 and (*E*) 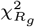 values for the HPS model (gray) and the optimized M1 (blue), M2 (orange) and M3 (green) models.

From the pool of optimized parameters, we selected the λ set which resulted in the largest Spearman’s correlation coefficient (*ρ* = 0.78) between simulated and experimental *R_g_* values for the A1 LCD variants. We base this final selection of the optimal λ set on the Spearman’s correlation coefficient of the A1 LCD variants because we expect that capturing the experimental ranking in chain compaction will result in accurate predictions of the relative LLPS propensities (15,16,20,58,59). Further, the systematic mutagenesis studies enable us to more clearly decouple the parameters for Y-vs-F and R-vs-K (15, 16). We note that while this selection uses only the A1 LCD variants, all three parameter sets result in good agreement with the full PRE and *R_g_* data set (Fig. 3*A*).

The selected model, referred to as M1 hereafter, is the starting point for two consecutive optimization cycles (Fig. S1*B*) which were performed with a lower weight for the prior (*θ* = 0.05), yielding a new pool of optimized parameters (squares in Fig. 3*A*) and model M2 (largest *ρ* = 0.75). To generate a third model, we further decreased the confidence parameter to *θ* = 0.02 and performed an additional optimization run starting from M2 (Fig. S1*C*). From the collected optimal parameters (triangles in Fig. 3*A*), we selected M3 (largest *ρ* = 0.73). As shown in Fig. 3*B*, the optimal λ values collected through the four independent optimization runs (Fig. S1*A*–*D*) are weakly intercorrelated. The covariance values range between −0.015 and 0.015 for most amino acids, with the exception of the standard deviations of N, C, T, M, W, and I. C, M, W, and I are among the least frequent amino acids in the training set (Fig. S3) and, unsurprisingly, we observe the largest covariance values for C-W (0.017), C-M (−0.02) and C-I (−0.016). Fig. 3*C* shows that M1–3 fall within two standard deviations (SDs) above and below the mean of the λ values yielding 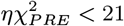 and 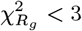 (gray shaded area). Despite their differences, M1–3 fit the training data equally accurately and result in an improvement in 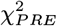 and 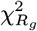 of ~30% and ~95% with respect to the HPS model, respectively (Fig. 3*D* and *E*).

Notably, the optimization procedure captures the sequence dependence of the chain dimensions (Fig. 4) and results in accurate predictions of intramolecular PRE data for both highly soluble IDPs and proteins that more readily phase separate (Fig. S4*B*–*D* and Fig. S5–S10), as well as in radii of gyration with relative errors −14% < Δ*R_g_* / *R_g,exp_* < 12% (Fig. S4*E* and *F*). Besides reproducing the experimental *R_g_* values for the longer chains with high accuracy, the optimized models also capture the differences in *R_g_* and scaling exponents, *v*, for the variants of A1 LCD (Fig. 4*B* and S11). The lower Pearson’s correlation coefficients observed for *ν*, compared to the corresponding *R_g_* data, may originate from the different models used to infer *ν* from SAXS experiments and simulation data, i.e., respectively, the molecular form factor method (16, 52) and least-squares fit to long intramolecular pairwise distances, *R_ij_*, vs |*i* − *j*| > 10 (60) (Fig. S12).

**Fig. 4.**
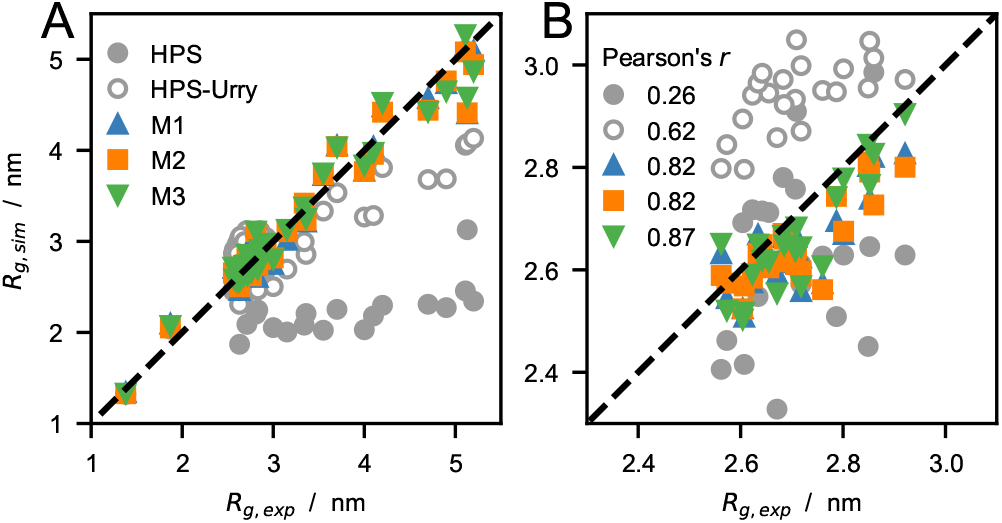
(*A*) Comparison between experimental and predicted radii of gyration (Tab. S1), *R_g_*, for the HPS, HPS-Urry, and M1–3 models. (*B*) Zoom-in on the *R_g_* values of the A1 LCD variants, with Pearson’s *r* coefficients for this subset of the training data reported in the legend.

To assess the impact of phase separating proteins on the optimized models, we perform an optimization run wherein the A1 LCD variants are removed from the training set. The major difference between the resulting optimal λ set and models M1– 3 is the considerably smaller values for R and Y residues (Fig. S2*C*). Indeed, the large λ values for R and Y residues in M1–3 relative to the HPS, AVG and HPS-Urry models, is a striking feature which resonates with previous experimental findings pointing to the important role of R and Y residues in driving LLPS (8,14–16,22,61,62).

To identify the hydrophobicity scales which most closely resemble M1–3, we construct a dendrogram (Fig. S13) complementing the 87 scales retained from the set by Simm et al. (39) with the Urry, Kapcha-Rossky and M1–3 scales, and using average linkage-based hierarchical clustering and Euclidean distances as the metric. This analysis reveals that the hydrophobicity scales by Urry et al. (29), Bishop et al. (42), Wimley and White (63) and the membrane protein surrounding hydrophobicity scale by Ponnuswamy and Gromiha (64) are those with greatest similarity to M1–3. These scales, which are characterized by a λ value for the R residue above the 80% quantile, are possibly the best of the unmodified scales for the properties that we optimized M1–3 to reproduce.

### Testing Protein-Protein Interactions

To test whether the parameters trained on single-chain conformational properties are transferable to protein-protein interactions, we compared experimental intermolecular PRE rates, Γ_2_, of FUS LCD and A2 LCD (22, 23) with predictions from two-chain simulations of the M1–3 models performed at the same conditions as the reference experiments. Intermolecular Γ_2_ values were obtained from solutions of spin-labeled ^14^N protein and ^15^N protein without a spin-label in equimolar amount and report on the transient interactions between a paramagnetic nitroxide probe attached to a cysteine residue of the spin-labeled chain and all the amide protons of the ^15^N-labeled chain. We carried out the calculation of the PRE rates using DEER-PREdict (34), assuming an effective correlation time of the spin label, *τ_t_*, of 100 ps and fitting an overall molecular correlation time, *τ_c_*, within the interval 1 ≤ *τ_c_* ≤ 20 ns. In agreement with experiments, Γ_2_ values predicted by the M1–3 models are characterized by no distinctive peaks along the protein sequence (Fig. 5*A*–*E*), which is consistent with transient and non-specific protein–protein interactions. Notably, while PRE rates for FUS LCD are of the same magnitude for all spin-labeled sites, the A2 LCD presents larger Γ_2_ values for S99C than for S143C indicating that the tyrosine-rich aggregation-prone region (residues 84–107) is involved in more frequent intermolecular contacts with the entire sequence. The discrepancy between predicted and experimental intermolecular PRE data, 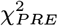, varies significantly as a function of *τ_c_* (Fig. 5*F*–*G*). For both FUS LCD and A2 LCD, the optimal *τ_c_* is larger for M1 than for M3, which suggests that the latter has more attractive intermolecular interactions. While for M1 the minimum of 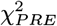 is at *τ_c_* = 17 ns for both proteins, for M3 the optimal *τ_c_* value is ~ 8 ns smaller for FUS LCD than A2 LCD. Although the accuracy of *τ_c_* is difficult to assess in the case of transiently interacting IDPs, this large difference in *τ_c_* (Fig. 5) suggests that the protein-protein interactions predicted for FUS LCD by M3 may be overly attractive.

**Fig. 5.**
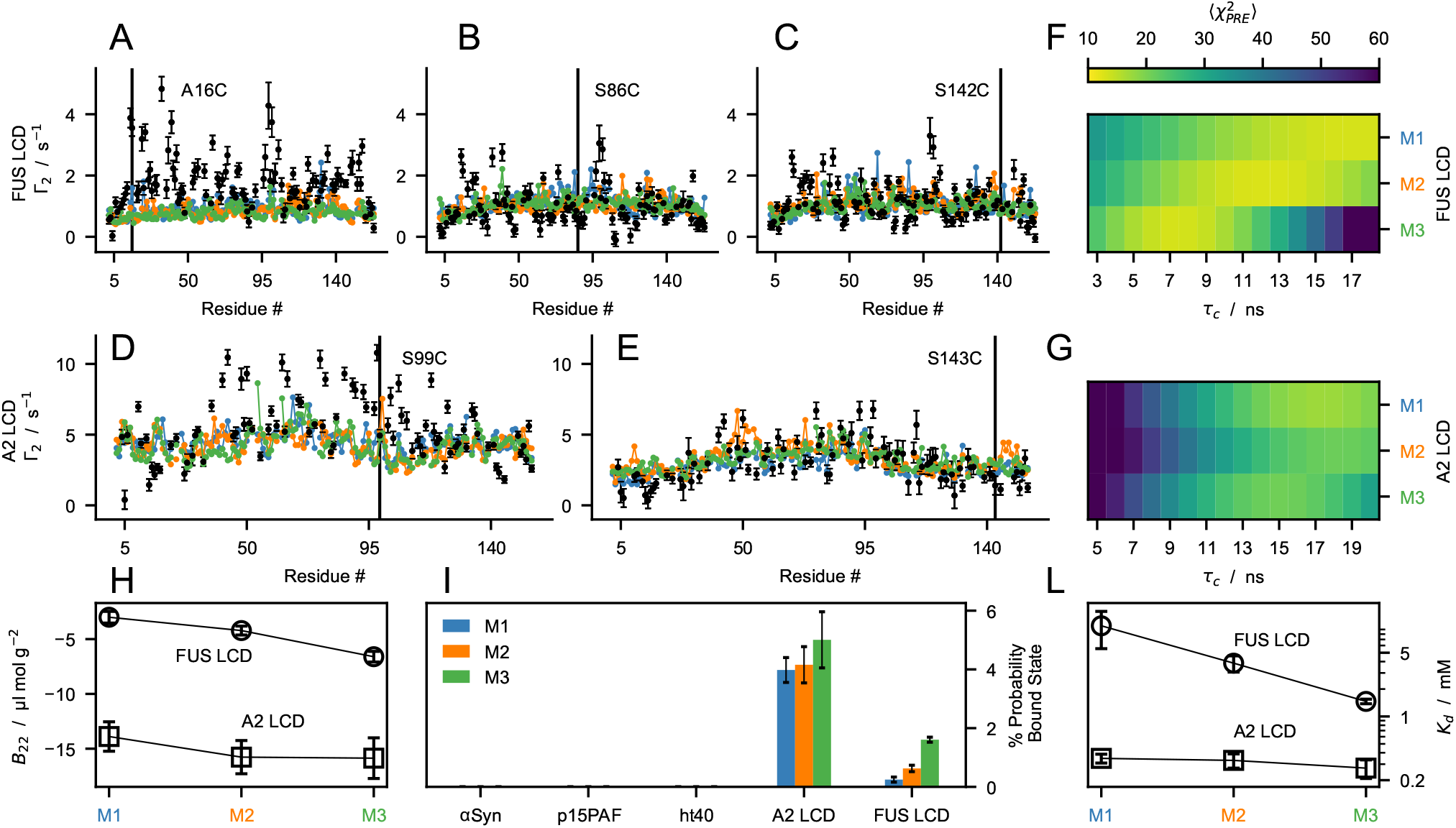
Testing the M1–3 models using experimental findings on protein-protein interactions. (*A–E*) Comparison between experimental (black) intermolecular PRE rates (Tab. S3) and predictions from the M1 (blue), M2 (orange) and M3 (green) models for FUS LCD (*A*–*C*) and A2 LCD (*D*–*E*) calculated using the best-fit correlation time, *τ_c_*. (*F*–*G*) Discrepancy between calculated and experimental intermolecular PRE rates 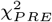 as a function of *τ_c_*. (*H*) Second virial coefficients, *B*_22_, of FUS LCD (circles) and A2 LCD (squares) calculated from two-chain simulations of the M1–3 models. Error bars are SEMs estimated by bootstrapping 1,000 times 40 *B*_22_ values calculated from trajectory blocks of 875 ns. (*I*) Probability of the bound state estimated from protein-protein interaction energies in two-chain simulations of the M1–3 models. (*L*) Dissociation constants, *K_d_*, of FUS LCD (circles) and A2 LCD (squares) calculated from two-chain simulations of the M1–3 models. For *p_B_* and *K_d_*, error bars are SDs of ten simulation replicas. Lines in *H* and *L* are guides to the eye.

To quantify protein-protein interactions with the optimized models, we calculated second virial coefficients, *B*_22_, from two-chain simulations (*SI Appendix*). The net interactions are attractive for both the sequences (*B*_22_ < 0), and considerably stronger for A2 LCD than for FUS LCD. As expected from the λ values and amino acid compositions, M3 presents the most negative *B*_22_ values (large λ values for Q, G and P), followed by M2 and M1 (Fig. 5*I*).

To test whether predictions of protein self-association by M1–3 are sequence dependent, we compared the probability of finding proteins in the bound dimeric state, *p_B_*, in simulations of *α*-Synuclein, p15PAF, full length tau (ht40), A2 LCD and FUS LCD performed at the solution conditions of the reference experimental data (37,50,65) (*SI Appendix*). In agreement with experimental findings, we find that the highly soluble *α*-Synuclein, p15PAF and ht40 proteins do not self-associate substantially in our simulations, whereas A2 LCD and FUS LCD have *p_B_* ~4% and ~1%, respectively. We further estimated the dissociation constants of A2 LCD and FUS LCD using *K_d_* = (1–*p_B_*)^2^/(*N_*Ap_B_*_V*) and *K_d_* = 1/(*N_Ap_B__*(*V*−*B*_22_)) self-consistently (66), where *N_A_* is Avogadro’s number (*SI Appendix*, Fig. *5L* and S14).

### Testing LLPS propensities

To test the ability of the models to capture the sequence-dependence of LLPS propensity, we performed multi-chain simulations in a slab geometry and calculated protein concentrations of the coexisting condensate, *c_con_*, and dilute phase, *c_sat_*. We compared our simulation results to an extensive set of sequences which have been shown to undergo LLPS below an upper critical solution temperature (UCST), namely FUS LCD (23,35,36), A2 LCD (22, 24), the NtoS variant of A2 LCD (24), LAF-1 RGG domain (11,67–69), as well as variants of A1 LCD (15, 16) and Ddx4 LCD (8,10,13). From simulations of the optimized models at 37°C, we observed that, for a number of sequences in the test set, the predicted *c_sat_* values are too low to allow for converged estimates from μs-timescale trajectories (Fig. S15). Conversely, the least LLPS-prone variants of Ddx4 LCD yielded one-phase systems when simulated at 37°C using HPS-Urry and M1–3 models. Thus, to be able to estimate converged *c_sat_* values (Fig. S16, S17 and S18), simulations were carried out at 50°C, except for the HPS-Urry model which we simulated at 24°C (Tab. S4). The FtoA and RtoA variants of Ddx4 LCD were also simulated at 24°C using the M1–3 models as in simulations of the same systems at 50°C we only observed a single phase.

Simulations using M1 at 50°C most closely recapitulate the experimental trend in *c_sat_* across the diverse sequences (Fig. 6*A*, *D* and *G*) and reproduce the reference *c_con_* and *c_sat_* values measured at room temperature. Conversely, HPS over-estimates the relative LLPS propensity of FUS LCD, whereas simulations using HPS-Urry at 24°C show deviations of about an order of magnitude from the reference *c_sat_* values for A2 LCD, Ddx4 LCD, A1 LCD and FUS LCD. Regarding the LAF-1 RGG domain, all of the models overestimate by at least a factor of ~5 the experimental *c_con_* (68, 69), whereas M1 reproduces within a factor of ~2 the experimental *c_sat_* value from temperature-dependent turbidity measurements (11), both for the WT and for variants with randomly shuffled sequence (LAF-1 shuf) and without residues 21–30 (LAF-1 Δ21–30) (Fig. S19). Although M1–3 fit the training data equally well, the prediction of LLPS propensities for the diverse sequences in Fig. 6*A* and *D* differ considerably, with Pearson’s correlation coefficients between simulation and experimental log_10_(*c_sat_*) values ranging from 0.67 for M1 to 0.14 for M3 (Fig. 6*G*). The discrepancy is particularly evident for the Ddx4 LCD and FUS LCD which are rich in N and Q residues, respectively, i.e., the residues for which the M1 and M3 λ sets differ the most.

**Fig. 6.**
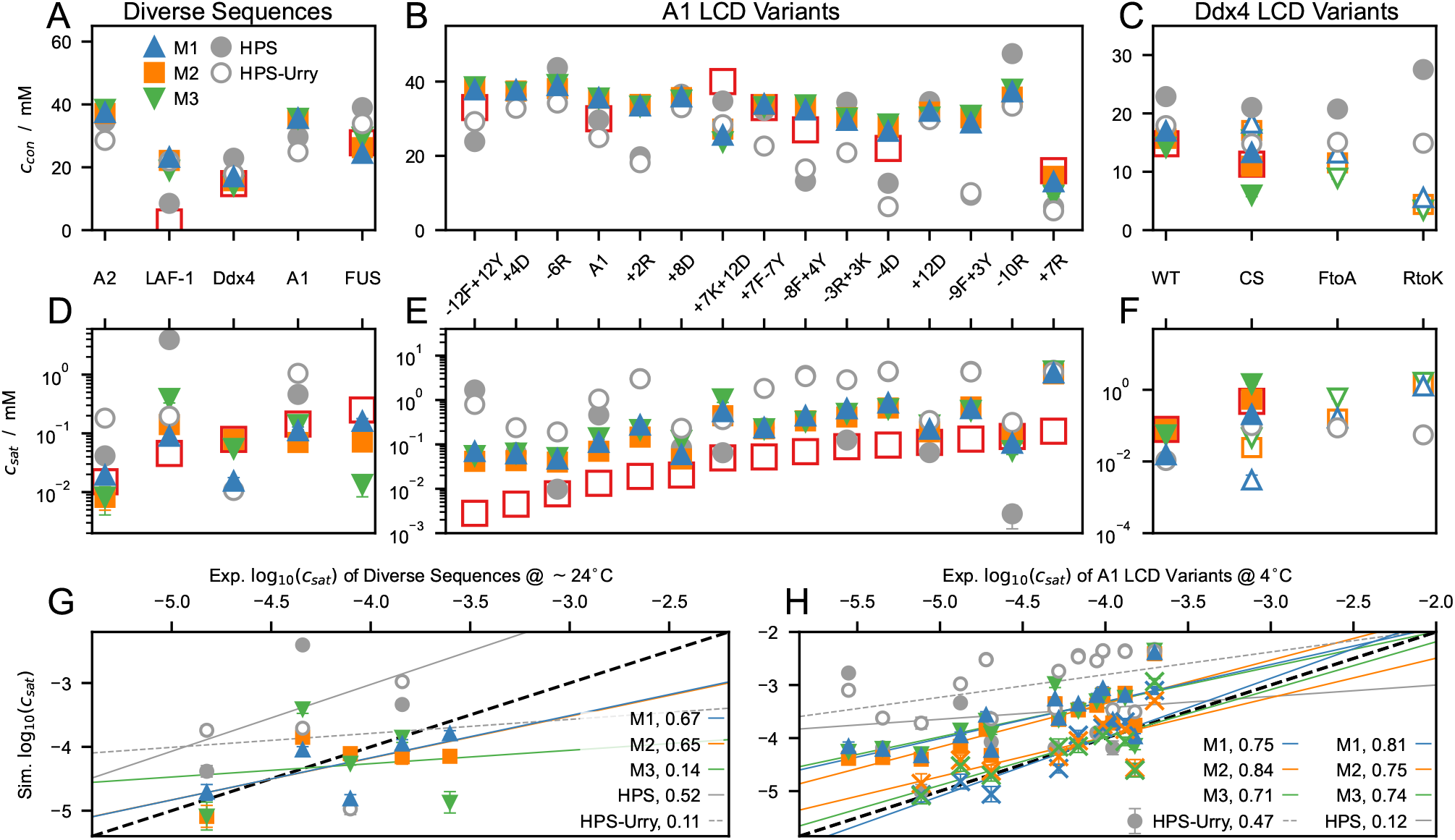
Protein concentrations in the condensate (*A*–*C*) and in the dilute phase (*D*–*F*) from slab simulations of the M1–3, HPS and HPS-Urry models performed at 50°C (closed symbols), 37°C (crosses in (*H*)) and 24°C (open symbols). Red open squares show experimental measurements at 24 C (*A, D, C* and *F*) and 4°C (*B* and *E*). (*G* and *H*) Correlation between log_10_(*c_sat_*/M) from simulations and experiments for diverse sequences (*G*) and A1 LCD variants (*H*). Solid lines show linear fits to the simulation data at 50°C. Dashed lines show linear fits to the HPS-Urry data at 24°C (*G* and *H*) and to the M1–3 data at 37°C (*H*). Values reported in the legends are Pearson’s correlation coefficients. Error bars are SEMs of averages over blocks of 0.3 μs. We note that the correlation coefficients reported in *G* are associated with a substantial uncertainty as they are calculated over only three (HPS), four (HPS-Urry) and five points (M1–3).

We further test our predictions against 15 variants of A1 LCD (Fig. 6*B* and *E*). These include aromatic and charge variants, which were designed to decipher the role on the driving forces for phase separation of Y vs F residues and of R, D, E and K residues, respectively (16). The nomenclature, ±*N_X_*X±*N_Z_*Z, denotes increase or decrease in the number of residues of type X and Z with respect to the WT, which is achieved by mutations to or from G and S residues while maintaining a constant G/S ratio. M1–3 are found to be equally accurate, and present a considerable improvement over previous models with respect to their ability to recapitulate the trends in LLPS propensity for the aromatic and charged variants of A1 LCD. Since M1–3 were selected based on their performance in predicting the experimental ranking for the *R_g_* values of 21 A1 LCD variants (Tab. S1), this result supports our model development strategy. For M1–3, Pearson’s correlation coefficients exceed 0.7 between log_10_(*c_sat_*) values measured at 4°C (16) and simulation predictions at both 50°C and 37°C (Fig. *6H*). Moreover, *c_sat_* values from simulations at 37°C are in agreement with the reference *c_sat_* values at 4°C (Fig. *6H* and S15). As we observed for the diverse sequences, quantitative agreement with the experimental *c_sat_* values is achieved by carrying out simulations of the M1 model at a temperature systematically larger by ~30°C than the experimental conditions. In addition to the lack of temperature dependence of the hydropathy parameters (70), the inconsistency between the temperature dependence of chain compaction and phase separation might be attributed to more general aspects of the model. For instance, the significant decrease in the number of interaction sites upon coarse-graining at the amino-acid level, and the resulting reduction in configurational entropy (71, 72), which may promote LLPS by lowering the entropic penalty associated with partitioning a chain from the dilute solution to the condensate.

M1–3 reproduce the experimental ranking for LLPS propensity of the Ddx4 LCD variants, i.e. WT≫CS>FtoA≳RtoK (Fig. 6*C* and *F*) and, for all the variants, M1 and M3 consistently display the highest and lowest LLPS propensities, respectively. Simulations at 50°C using M2 are in quantitative agreement with the experimental *c_sat_* values (13) for both WT and the CS variant, which has the same net charge and amino acid composition as the WT but a more uniform charge distribution along the sequence. Moreover, as observed experimentally (13), M1–3 predict a single phase for the RtoK variant at 24°C. As previously shown by Das et al. (25), the HPS model predicts a considerable increase in LLPS propensity upon replacement of all 24 R residues in the Ddx4 LCD with K (RtoK variant; Fig. 6*C*), in apparent contrast to experimental observations (10, 13). Interestingly, augmenting the HPS model with stronger cation-*π* interactions for R-aromatic than for K-aromatic pairs (25) has been shown to be insufficient to capture the lower LLPS propensity of the RtoK variant compared to WT. On the other hand, our data for the M1–3 and HPS-Urry models indicates that making all the interactions involving R more favourable results in more accurate predictions. In fact, a large λ value for R may better mimic its relatively unfavorable free energy of hydration (19) as well as the occurrence of R-aromatic cation-*π* interactions, R-R *π*-stacking and R-D/E bidentate H-bonding (10,17,18,73). Compared to the Kapcha-Rossky scale, it is noteworthy that the increase in the λ values of R, Y and G in M1–3 is accompanied by an overall decrease in the average λ value. Hence, the optimization procedure led to the enhancement of specific attractive forces while maintaining a balance between electrostatic and non-electrostatic interactions (25), which reveals itself, for example, in the ability of M1–3 to recapitulate the lower LLPS propensity of the CS variant with respect to Ddx4 LCD WT.

The M1 and M2 parameter sets differ mainly for the λ value of the N residue (Fig. 3*C*) and perform equally well against the test set (Fig. 6). Therefore, we further test the ability of M1 and M2 to predict the LLPS propensity of the NtoS variant of A2 LCD with respect to the wild type. Only the M1 model, which has λ values for N and S of similar magnitude correctly predicts approximately the same LLPS propensity for variant and WT (Fig. S20), in agreement with experiments (24).

### Correlating single-chain properties and phase separation

Motivated by recent experiments on the A1 LCD (15, 16), we perform a detailed analysis of the coupling between chain compaction and phase behaviour of the A1 LCD variants. In agreement with previous observations (16), the log_10_(*c_sat_*) values for the aromatic variants show a linear relationship with the scaling exponent, *ν_sim_*, whereas changes in the number of charged residues (charge variants) result in significant deviations from the lines of best fit (Fig. 7*A*–*C*). Following the approach of Bremer, Farag et al. (16), we plot the residuals for the charge variants with respect to the lines of best fit as a function of the net charge per residue (NCPR) (Fig. 7*D*–*F*). The results for M1 and M2 show the V-shaped profile observed for the experimental data (16), and support the suggestion that mean-field electrostatic repulsion between the net charge of the proteins is responsible for breaking the coupling between chain compaction and LLPS propensity (16). In agreement with experimental data (16), we observe that for M1 and M2 the driving forces for LLPS are maximal for small positive values of NCPR (~0.02).

**Fig. 7.**
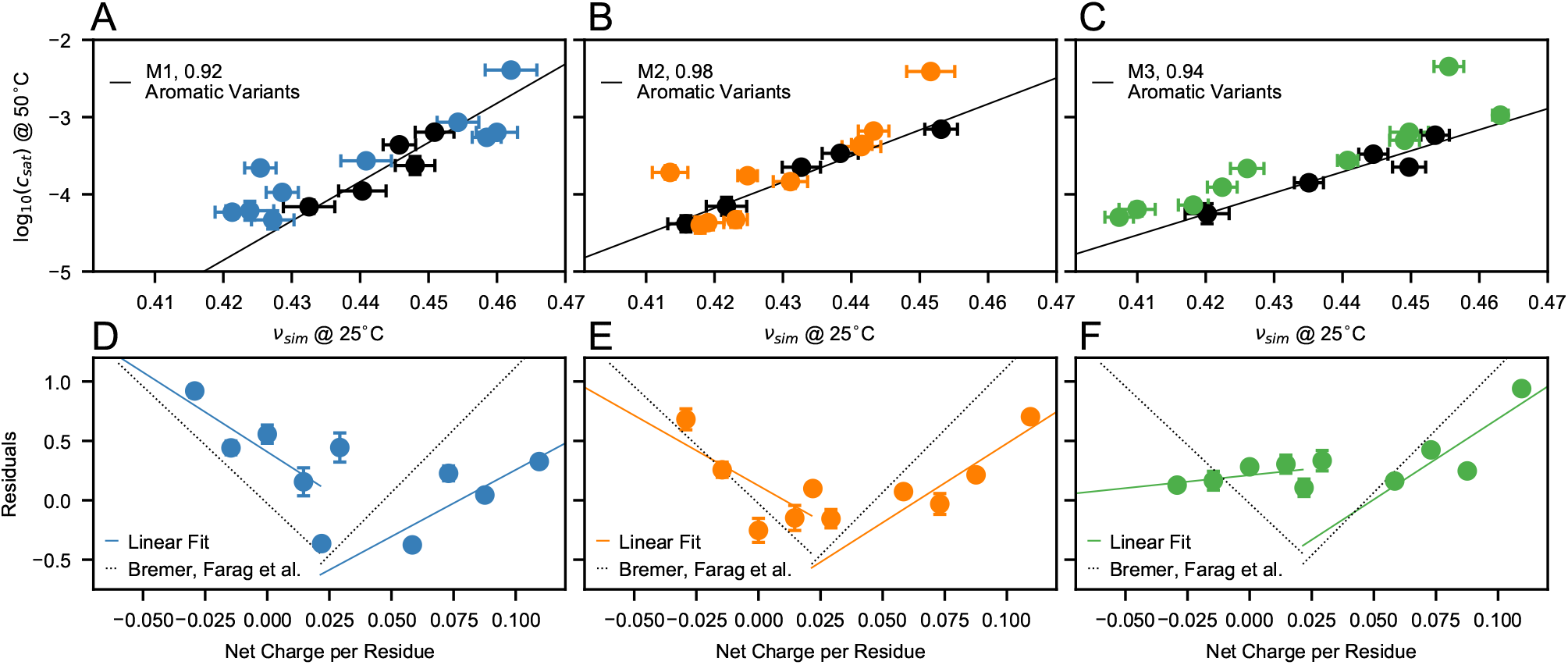
Correlation between chain compaction and LLPS propensity for aromatic and charge variants of A1 LCD. (*A*–*C*) log_10_(*c_sat_*) vs *ν_sim_* for A1 LCD variants from simulations performed using the M1 (*A*), M2 (*B*) and M3 (*C*) models. Black and colored circles indicate aromatic and charge variants, respectively. Black lines are linear fits to the aromatic variants. (*D*–*F*) Residuals from the linear fits of panels *A*–*C* for the charge variants of A1 LCD as a function of the net charge per residue. Values reported in the legends are Pearson’s correlation coefficients. Error bars of log_10_(*c_sat_*) values are SEMs of averages over blocks of 0.3 μs. Error bars of *ν_sim_* are SDs from fits to *R_ij_* = *R*_0_|*i*−*j*|^*ν*_*sim*_^ in the long-distance region, |*i*−*j*| > 10. Solid lines are linear fits to the data. Dotted lines in *D*–*F* are lines of best fit to the experimental data by Bremer, Farag et al. (16).

The dependence of LLPS on NCPR is clarified by comparing the residual non-electrostatic energy maps of +8D (NCPR=0), +4D (NCPR≈0.03) and −4D (NCPR≈0.09) with respect to the wild type of A1 LCD (NCPR≈0.06) (Fig. S21 and S22). While in the case of NCPR=0 the residual interaction patterns within the isolated chain and between chains in the condensate largely overlap, the energy baselines are clearly down- and up-shifted for NCPR≈0.03 and NCPR≈0.09, respectively (Fig. S21*G*–*I* and S22*G*–*I*). Although the interaction patterns are still dominated by the stickers, deviations of the NCPR from —0.02 result in electrostatic mean-field repulsive interactions that disfavor LLPS. The LLPS-promoting effect of small positive NCPR values finds explanation in the amphiphilic character of the R side chains (18) which compensate for the repulsion introduced by the excess positive charge by allowing for favorable interactions with both Y and negatively charged residues.

As opposed to M1–2, the readily phase-separating M3 model shows a weaker dependence on NCPR, especially for variants of net negative charge. This suggests that the experimental observations regarding the coupling between conformational and phase behaviour of A1 LCD stem from a well-defined balance between mean-field repulsion and sticker-driven LLPS which can be offset by an overall moderate increase of 3–4% in the λ values of the residues present in A1 LCD.

### Comparing intra- and inter-molecular interactions

After establishing the ability of model M1 to accurately predict trends in LLPS propensity for diverse sequences, we analyze the non-electrostatic residue-residue energies for FUS LCD and A2 LCD within a single chain, as well as between pairs of chains in the dilute regime and in condensates. We find a striking similarity between intra- and intermolecular interaction patterns for both proteins (Fig. 8), consistent with a mostly uniform distribution of stickers along the linear sequence (Fig. 8*G* and *H*) (15, 74). Notably, besides the aromatic F and Y residues, the analysis also identifies an M residue and four R residues as stickers in FUS LCD and A2 LCD, respectively. Therefore, the parameter-learning procedure presented herein corroborates the important role of R as a sequence dependent sticker (16), whereby the large λ value for R in models M1–3 presumably reflects the ability of the amphiphilic guanidinium moiety to engage in H-bonding, as well as *π* stacking and charge-*π* interactions (18). Further, in the dilute regime, the intra- and intermolecular interactions are weaker in the N- and C-terminal regions than for the rest of the chain, as evident from the upturning baselines of the 1D interaction energy projections. This result is consistent with the faster local motions of the terminal residues inferred from ^15^N NMR relaxation data for both unfolded proteins (75) and a number of phase separating IDPs (15,22,23). We also find that the aggregation-prone Y-rich region of A2 LCD (residues 84–107) interacts with the entire polypeptide chain (Fig. 8*D*–*F*) and thus likely drives chain compaction, self-association as well as LLPS. Finally, in line with previous observations from theory, simulations and experiments (16,76,77), we observe that the polypeptide chains of A1 LCD, A2 LCD and FUS LCD are more expanded in the condensed phase than in the dilute phase (Fig. S23). In particular, we find that the scaling exponents of the LCDs increase towards *ν* = 0.5 in the condensed phase, and that differences in compaction between wild-type and charge variants of A1 LCD are greater in the dilute than in the condensed phase (Fig. S23).

**Fig. 8.**
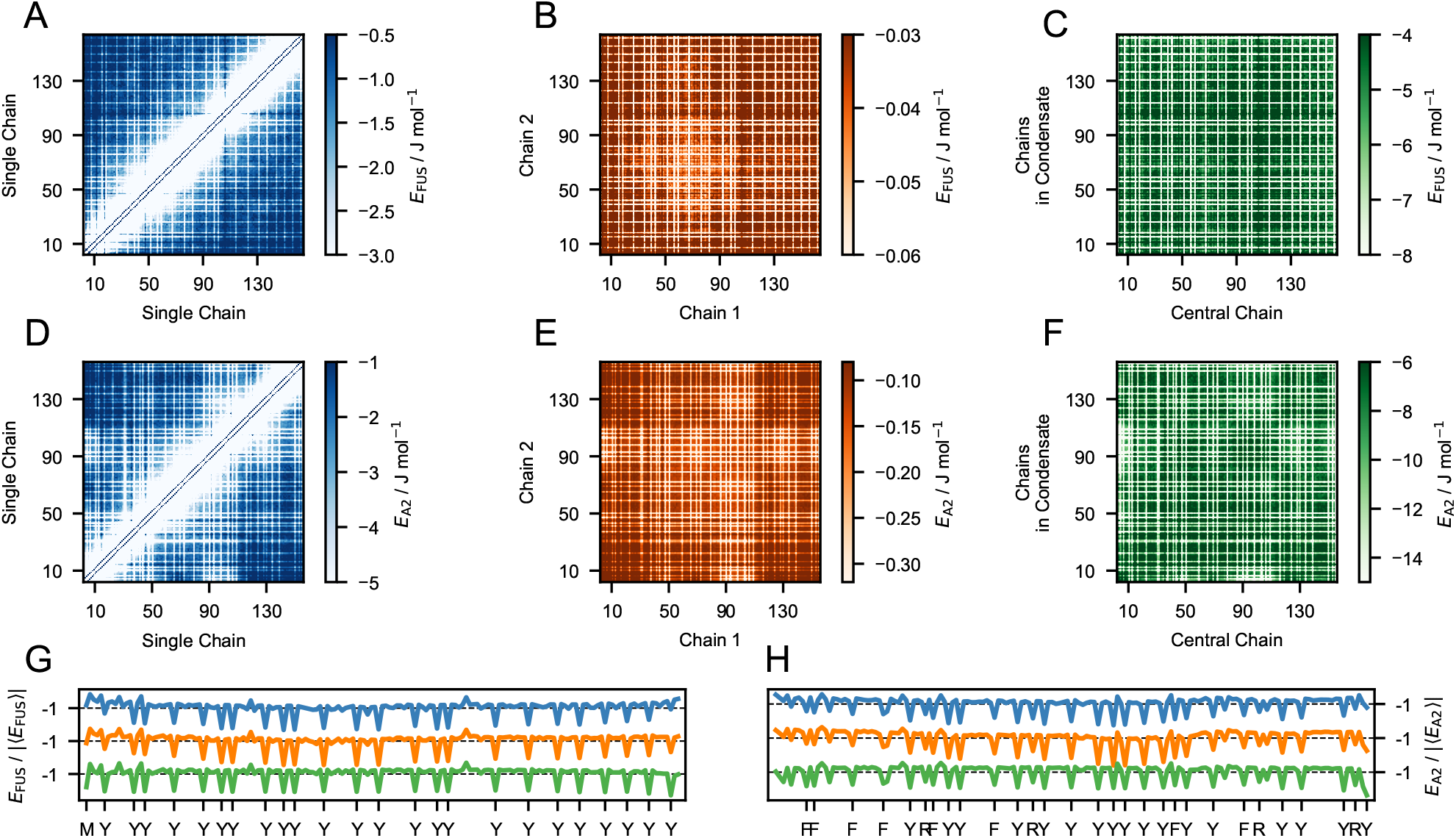
Comparing residue-residue interactions in dilute solution and in the condensate. Energy maps from simulations of the M1 model of FUS LCD (*A–C*) and A2 LCD (*D–F*) calculated using non-electrostatic interaction energies. (*G–H*) 1D projections of the energy maps for FUS LCD (*G*) and A2 LCD (*H*), normalized by the absolute average interaction energy |〈*E*〉| and shifted vertically for clarity. Colors indicate that the energies were calculated within a single chain at infinite dilution (blue), between two chains in the dilute regime (orange) and between a chain located at the center of a condensate and the surrounding chains (green).

## Conclusions

In this work we implement and validate an automated procedure to develop an accurate model of the LLPS of IDPs based on experimental data reporting on single-chain conformational properties. We show that this strategy succeeds, in agreement with the previously observed coupling between chain compaction and propensity for phase separation (15,20,58,59), but also appears to recapitulate the recent discovery that charge effects may break this relationship (16). Our work differs from related previous studies (28,30,33,78) in several ways including the size of the data set used for optimization, the use of both NMR PREs and *R_g_* values, and the introduction of a prior for the λ values. Moreover, by carrying out model optimizations with and without the A1 LCD variants, we show that the presence of phase-separating IDPs in the training set helps the parameter-learning procedure to capture the role of Y and R residues as stickers. The accuracy and general applicability of our model can be tested further by future experiments on systems that were not used for training or testing. We also note that our automated, Bayesian optimization approach makes it relatively straightforward to continue to develop and improve the model as additional data becomes available.

Simulations performed using the model optimized herein reveal that, at least for sequences characterized by a relatively uniform distribution of stickers, residue-residue interactions determining chain compaction also drive self-association and LLPS. Moreover, we show that the experimentally-observed dependency of LLPS on protein net charge appears to be captured by salt-screened electrostatic repulsion, even when assuming a uniform dielectric constant throughout the two-phase system.

We have here shown how our model may be used to help elucidate the residues that are important for LLPS of IDPs with UCST behaviour. Further, we suggest the model could be applied to study the influence of disease-associated mutations on the material properties of protein self-coacervates (79, 80), the LLPS of protein mixtures as a function of composition, and the partitioning of proteins that do not readily undergo phase separation alone into condensates formed by other proteins (81, 82). Finally, owing to the generalized parameter-learning approach, the model could readily be refined as new experimental data are collected and it should be possible to extend it to account for specific pairwise interactions such as cation-*π* interactions (25), PTMs (83), the salting-out effect (84) and the temperature dependence of solvent mediated interactions (70).

## Materials and Methods

We use the C*α*-based model proposed by Dignon et al. (21) augmented with extra charges for the termini and a temperature-dependent treatment for dielectric constant of water (*SI Appendix*). Langevin dynamics simulations are conducted using HOOMD-blue v2.9 (85) in the *NVT* ensemble using the Langevin thermostat with a time step of 5 fs and friction coefficient 0.01 ps^−1^ (*SI Appendix*). Additionally, 100- and 300-chain simulations of LAF-1 RGG domain are also performed using openMM v7.5 (86) (Fig. S20).

### Bayesian Parameter-Learning Procedure

The λ values are optimized using a Bayesian parameter-learning procedure (30,87,88). The training set consists of the experimental *R_g_* values of 42 IDPs (Tab. S1) and the intramolecular PRE data of six proteins (Tab. S2) (16,22,23,37,44–57). To guide the optimization within physically reasonable parameters and to avoid over-fitting the training set, we introduce a regularization term which penalizes deviations of the λ values from the probability distribution, *P*(λ), which is the prior knowledge obtained from the statistical analysis of 87 hydrophobicity scales. The optimization procedure consists of the following steps (Fig. 2):

1. Single-chain CG simulation of the proteins of the training set (Tab. S1);
2. Conversion of CG simulations into all-atom trajectories using PULCHRA (89) of the proteins in Tab. S2 for the calculation of the PRE data;
3. Calculation of per-frame radii of gyration and PRE data. The PRE rates, Γ_2_, and intensity ratios, *I*_para_/*I*_dia_, are calculated using the rotamer library approach implemented in DEER- PREdict (34) with *τ_t_* = 100 ps and optimizing the correlation time, *τ_c_* ∈ [1, 10] ns, against the experimental data.
4. Random selection of six λ values which are nudged by random numbers picked from a normal distribution of standard deviation 0.05. The prior probability distribution, *P*(λ), sets the bounds of the parameter space: any λ_*i*_ for which *P*(λ_*i*_) = 0 is further nudged until *P*(λ_*i*_) = 0.
5. Calculation of the Boltzmann weights for the *i^th^* frame as *ω_i_* = exp −[*U*(***r_i_***,**λ**_*k*_) − *U*(***r_i_***,**λ_0_**)]/*k_B_T*, where *U*(***r_i_***, **λ**_***k***_) and *U*(***r_i_***, **λ**_0_) are the total Ashbaugh-Hatch energies of the *i^th^* frame for trial and initial λ values, respectively. If the effective fraction of frames,

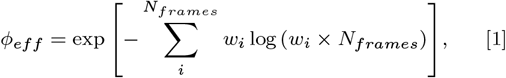

is below 30%, the trial **λ_*k*_** is discarded.
6. The per-frame radii of gyration and PRE observables are reweighted and the extent of agreement with the experimental data is estimated as

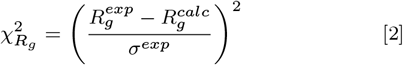

and

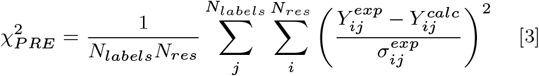

where 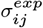 is the error on the experimental values, *Y* is either *I_para_*/*I_dia_* or Γ_2_, *N_labels_* is the number of spin-labeled mutants and *N_res_* is the number of measured residues;
7. Following the Metropolis criterion (90), the *k^th^* set of λ values is accepted with probability:

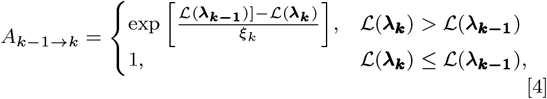

where the control parameter, *ξ_k_*, scales with the number of iterations as *ξ* = *ξ*_0_ × 0.99^*k*^. 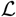 is the cost function

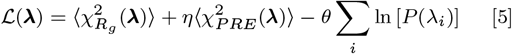

where 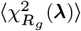 and 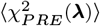 are averages over the proteins in the training sets. *θ* and *η* are hyperparameters of the optimization procedure. *θ* determines the trade-off between between over- and under-fitting the training set whereas *η* sets the relative weight of the PRE data with respect to the radii of gyration.

Steps 4–7 are iterated until *ξ* < 10^−15^, when the reweighting cycle is interrupted and new CG simulation carried out with the trained λ values. A complete parameter-learning procedure consists of two reweighting cycles starting from *ξ*_0_ = 2 followed by three cycles starting from *ξ*_0_ = 0.1. The threshold on *φ_eff_* results in average absolute differences between *χ*^2^ values estimated from reweighting and calculated from trajectories performed with the corresponding parameters of ~1.8 and ~0.8 for 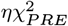 and 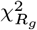, respectively (Fig. S24).

### Data deposition

Datasets, code and Jupyter Notebooks for reproducing our analyses are available on GitHub at github.com/KULL-Centre/papers/tree/main/2021/CG-IDPs-Tesei-et-al and on Zenodo, DOI: 10.5281/zenodo.5005953.

## ACKNOWLEDGMENTS

We thank Veronica Ryan and Nicolas L. Fawzi for sharing the PRE data for FUS LCD, FUS12E LCD and A2 LCD as well as Robert Konrat for sharing the intramolecular PRE data for Osteopontin. We thank Robert B. Best for sharing data on compaction of IDPs, Gregory L. Dignon and Jeetain Mittal for help setting up simulations with the HPS model, and Tanja Mittag, Massimiliano Bonomi and Benjamin Schuler for helpful discussions. We acknowledge funding from the BRAINSTRUC structural biology initiative from the Lundbeck Foundation (R155-2015-2666), and acknowledge access to computational resources from the ROBUST Resource for Biomolecular Simulations (supported by the Novo Nordisk Foundation; NNF18OC0032608) and Biocomputing Core Facility at the Department of Biology, University of Copenhagen. T.K.S. acknowledges support from the Novo Scholarship Programme 2021. This project has received funding from the European Union’s Horizon 2020 research and innovation programme under the Marie Skłodowska-Curie grant agreement No. 101025063.

