## Supporting Text, Figures and Tables for "Accurate model of liquid-liquid phase behaviour of intrinsically-disordered proteins from optimization of single-chain properties"

Datasets, code and Jupyter Notebooks for reproducing the presented analyses and plots can be accessed on GitHub at github.com/KULL-Centre/papers/tree/main/2021/CG-IDPs-Tesei-et-al and on Zenodo, DOI: 10.5281/zenodo.5005953.

### SI Materials and Methods.

***C$\alpha$-based model.*** In the simulations performed in this study, protein residues interact through the Ashbaugh-Hatch potential (1),

$$u_{AH}(r) = \begin{cases} u_{LJ} + \epsilon(1-\lambda), & r \le 2^{1/6}\sigma \\ \lambda u_{LJ}, & r > 2^{1/6}\sigma, \end{cases} \qquad [1]$$

where $\epsilon = 0.8368$ kJ mol$^{-1}$ and $u_{LJ}$ is the Lennard-Jones potential:

$$u_{LJ}(r) = 4\epsilon \left[ \left(\frac{\sigma}{r}\right)^{12} - \left(\frac{\sigma}{r}\right)^{6} \right], \qquad [2]$$

$\sigma$ and $\lambda$ are arithmetic averages of amino-acid specific parameters quantifying size (2) and hydrophobicity (3), respectively. For $\sigma$, we use the values calculated from van der Waals volumes by Kim and Hummer (2). The $\lambda$ values of the HPS model were obtained through min-max normalization of the hydrophobicity scale by Kapcha and Rossky (3, 4). The HPS-Urry model was instead derived from the scale proposed by Urry et al. and further optimized by shifting the $\lambda$ values by -0.08 (5, 6). As suggested in a recent study on the effect of N-acetylation on the LLPS of FUS LCD (7), the charged termini play a significant role in modulating LLPS. Therefore, in the present work, we implicitly model the N- and C-terminal amonium and carboxylate moieties by, respectively, increasing and decreasing by one unit the charge number of the terminal beads. Salt-screened electrostatic interactions are modeled via the Debye-Hückel potential,

$$u_{DH}(r) = \frac{q_i q_j e^2}{4\pi\epsilon_0\epsilon_r r} \exp(-r/D), \qquad [3]$$

where $q$ is the average amino acid charge number, $e$ is the elementary charge, $D = \sqrt{1/(8\pi B c_s)}$ is the Debye length of an electrolyte solution of ionic strength $c_s$ and $B(\epsilon_r)$ is the Bjerrum length. Electrostatic interactions are truncated and shifted at the cutoff, $r_{\mathrm{cut}} = 4$ nm. As opposed to the HPS model wherein $\epsilon_r = 80$, we model the implicit solvent through the temperature-dependent dielectric constant of water as expressed by the following empirical relationship (8):

$$\epsilon_r(T) = \frac{5321}{T} + 233.76 - 0.9297 \times T + 1.417 \times 10^{-3} \times T^2 - 8.292 \times 10^{-7} \times T^3, \qquad [4]$$

such that e.g. $\epsilon_r(310\text{ K}) = 77.7$ and $\epsilon_r(360\text{ K}) = 58.8$. The average charge of the histidine residues is estimated based on the Henderson–Hasselbalch equation using p$K_a = 6$ (9).
The amino acid beads are connected by harmonic potentials,

$$u_{bond}(r) = \frac{1}{2}k(r - r_0)^2, \qquad [5]$$

of force constant $k = 8033$ kJ mol$^{-1}$ nm$^{-2}$ and equilibrium distance $r_0 = 0.38$ nm.
We note that the HPS and HPS-Urry models that we used in this study differ from the original models proposed by Dignon et al. (4) and Regy et al. (6) by the modified charges of the terminal beads, the value of the Debye length and the temperature dependence of $\epsilon_r$.

***Molecular Simulations.*** Langevin dynamics simulations of individual chains are performed in a cubic box of side length 300 nm under periodic boundary conditions, starting from the fully extended conformation generated using PeptideBuilder (10). Non-bonded interactions between residues separated by one bond are excluded from the energy calculation. Systems are simulated for 155 ns discarding the intial 5 ns of the trajectories and saving 15,000 frames every 10 ps. To speed up the optimization procedure, simulations of long sequences are split into two (OPN, CoRNID, K23, K32) or three (PNt, K44) replicas. In two-chain simulations of $\alpha$-Synuclein, p15PAF, full length tau (ht40) and of the LCDs of FUS and A2, the side length of the cubic simulation box is initially 100–200 nm during a short simulation of 0.1–1 ns, after which it is shrunk to 25.5, 34, 48, 40.5 or 48 nm to match the experimental protein concentration of 200 (11), 85 (12), 30 (13), 50 (14) and 30 µM (15), respectively. For the the LCDs of FUS and A2, ten replicas of each system are further simulated for 3.5 µs each, saving every 30 ps. For $\alpha$-Synuclein, p15PAF and ht40 we simulated only two replicas of 2 µs each. In multi-chain simulations, 100 proteins are inserted in a cuboidal box of side lengths $[L_x, L_y, L_z] = [17, 17, 300]$ nm and $[L_x, L_y, L_z] = [15, 15, 150]$ nm for variants of Ddx4 LCD and the remainder sequences, respectively. Additionally, 300 chains of LAF-1 RGG domain are simulated in a cuboidal box of side lengths $[L_x, L_y, L_z] = [24, 24, 240]$ nm. No significant differences are observed with respect to the corresponding 100-chain simulation (Fig. S19*G* and *M*). The chains are initially aligned along the $z$-axis and placed in the mid-plane of the box at random $(x, y)$ positions so that they are more than 0.7 nm apart from each other. During a 200 ns equilibration run, the proteins are confined in a slab by two Gaussian walls located at $z_G = \pm 30$ nm and $z_G = \pm 10$ nm for variants of Ddx4 LCD and the remainder sequences, respectively:

$$u_G(z) = \begin{cases} \epsilon_G \exp\left[-\frac{1}{2}(z - z_G)^2\right], & z < z_{\text{cut}} \\ 0, & z \geq z_{\text{cut}}, \end{cases} \quad [6]$$

where $z$ is the distance from the mid-plane of the cuboidal simulation box, $z_{\text{cut}} = 4$ nm and $\epsilon_G = 10$ kJ mol$^{-1}$. Multi-chain systems are further simulated without Gaussian walls for at least 2 µs saving frames every 250 ps.

***Second virial coefficient.*** The second virial coefficient is calculated from two-chain simulations through the Meyer integral of the radial distribution function, $g(r)$, for the mass-center separation between the chains, $r$:

$$B_{22} = \int_0^\infty \mathrm{d}r \; 2\pi r^2 [g(r) - 1]. \quad [7]$$

To correct for the finite size of the simulated closed systems, we apply the empirical correction proposed by Ganguly and van der Vegt (16) to the $g(r)$

$$g^{correct}(r) = g(r) \frac{1 - V(r)/V_{cell}}{1 - V(r)/V_{cell} - \Delta N(r)}. \quad [8]$$

where $V_{cell}$ is the volume of the simulation cell, $V(r)$ is the volume of a sphere of radius $r$ and $\Delta N(r)$ is the excess number of proteins within $V(r)$,

$$\Delta N = \frac{2}{V} \int_0^r \mathrm{d}r \; 4\pi r^2 [g(r) - 1] \quad [9]$$

The nominator and denominator of Eq. 8 approximate the number of proteins in the remainder of the cell outside $V(r)$ in case of a uniform protein distribution and in the actual system, respectively.

***Dimer dissociation constant.*** The probability of observing a dimeric bound state, $p_B$, is estimated from two-chain simulations by applying a dual cutoff scheme to the time series of the intermolecular non-electrostatic interaction energy. To make the dual cutoff scheme applicable to protein sequences of any length, $N$, we normalize the intermolecular energy by $N \ln N$, i.e. the scaling law inferred from the average intermolecular non-electrostatic interaction energy, $\langle E \rangle$, calculated for polyvaline chains of 50, 100, 150, 200, 250, 300, 400, 500 and 600 residues modelled using the M1 model (Fig. S14*A*). We consider a binding event to occur when the normalized intermolecular energy, $e = E/(N \ln N)$, drops below the lower cutoff, $e_l$. The bound state is assumed to survive thereafter until it dissociates when $e$ exceeds the upper cutoff, $e_u$. After estimating $p_B$ as the fraction of frames where the chains are in the bound state, the dissociation constant can be estimated using $K_d = (1 - p_B)^2/(N_A p_B V)$ and $K'_d = 1/(N_A p_B (V - B_{22}))$, where $N_A$ is Avogadro's number. $K_d$ and $K'_d$ are evaluated for $-0.15 < e_l < -0.02$ kJ mol$^{-1}$ and $e_l + 0.01 < e_u < -0.01$ kJ mol$^{-1}$. Fig. S14*B* shows the values of $e_l$ and $e_u$ yielding $2(K_d - K'_d)/(K_d + K'_d) < 5\%$. From the region of maximum overlap between the data for FUS LCD and A2 LCD (white contour line in Fig. S14*B*), we identify the optimal pairs of [$e_l$,$e_u$] values as [-0.09,-0.03], [-0.08,-0.04], [-0.07,-0.04], [-0.07,-0.05] and [-0.06,-0.05] kJ mol$^{-1}$. Fig. S14*C* shows that the SDs of $p_B$ and $K_d$ estimated using the five optimal dual cutoffs are lower than or comparable to the SDs of ten simulation replicas calculated with [$e_l$,$e_u$]=[-0.07,-0.05] kJ mol$^{-1}$ and displayed in Fig. 5*I* and 5*L*.

***Phase Diagrams from multi-chain simulations.*** The slab is centered in the box by translating the particles along $z$ by a displacement $d_i$ which maximizes the correlation function,

$$K(d_i) = \int \mathrm{d}z \; \rho_0(z) \rho_i(z + d_i), \quad [10]$$

where $\rho_0(z)$ and $\rho_i(z)$ are the density profiles of the initial and $i^{th}$ frames, respectively. The equilibrium density profile, $\rho(z)$, is calculated from the average of $\rho_i(z + d_i)$ over the last 1 µs of the trajectory. The density of the dilute and protein-rich phases is estimated by fitting the semi-profiles in $z > 0$ and $z < 0$ to

$$\rho(z) = \frac{\rho_a + \rho_b}{2} + \frac{\rho_b - \rho_a}{2} \tanh\left(\frac{|z| - z_{DS}}{t}\right), \quad [11]$$

where $z_{DS}$ is the position of the dividing surface, $t$ is the thickness of the interfacial region whereas $\rho_a$ and $\rho_b$ are the densities of the protein-rich and dilute phases, respectively. The uncertainty of the density values is estimated as the standard deviation of the the average densities in the regions $|z| < z_{DS} - t/2$ and $|z| > z_{DS} + 4t$ nm calculated for blocks of 0.3 µs.

**Table S1. Solution conditions and gyration radii of proteins included in the training set. Shaded rows highlight the variants of A1 LCD.**

| Protein | $N$ | $R_g$ (nm) | $T$ (K) | $c_s$ (M) | pH | Ref. |
|---|---|---|---|---|---|---|
| Hst5 | 24 | $1.38 \pm 0.07$ | 293 | 0.15 | 7.5 | (17) |
| $(\text{Hst5})_2$ | 48 | $1.87 \pm 0.07$ | 298 | 0.15 | 7.0 | (18) |
| ACTR | 71 | $2.63 \pm 0.1$ | 278 | 0.2 | 7.4 | (19) |
| Sic1 | 92 | $3.00 \pm 0.4$ | 293 | 0.2 | 7.5 | (20) |
| SH4UD | 95 | $2.71 \pm 0.1$ | 293 | 0.2 | 8.0 | (21) |
| ColNT | 98 | $2.83 \pm 0.1$ | 277 | 0.4 | 7.6 | (22) |
| p15PAF | 111 | $2.81 \pm 0.1$ | 298 | 0.15 | 7.0 | (12) |
| hNL3cyt | 119 | $3.15 \pm 0.2$ | 293 | 0.3 | 8.5 | (23) |
| RNaseA | 124 | $3.36 \pm 0.1$ | 298 | 0.15 | 7.5 | (24) |
| A1 | 137 | $2.76 \pm 0.05$ | 298 | 0.15 | 7.0 | (25) |
| -10R | 137 | $2.67 \pm 0.05$ | 298 | 0.15 | 7.0 | (25) |
| -6R | 137 | $2.57 \pm 0.05$ | 298 | 0.15 | 7.0 | (25) |
| +2R | 137 | $2.62 \pm 0.05$ | 298 | 0.15 | 7.0 | (25) |
| +7R | 137 | $2.71 \pm 0.05$ | 298 | 0.15 | 7.0 | (25) |
| -3R+3K | 137 | $2.63 \pm 0.05$ | 298 | 0.15 | 7.0 | (25) |
| -6R+6K | 137 | $2.79 \pm 0.05$ | 298 | 0.15 | 7.0 | (25) |
| -10R+10K | 137 | $2.85 \pm 0.05$ | 298 | 0.15 | 7.0 | (25) |
| +12D | 137 | $2.80 \pm 0.05$ | 298 | 0.15 | 7.0 | (25) |
| +4D | 137 | $2.72 \pm 0.05$ | 298 | 0.15 | 7.0 | (25) |
| +8D | 137 | $2.69 \pm 0.05$ | 298 | 0.15 | 7.0 | (25) |
| -9F+3Y | 137 | $2.68 \pm 0.05$ | 298 | 0.15 | 7.0 | (25) |
| +12E | 137 | $2.85 \pm 0.05$ | 298 | 0.15 | 7.0 | (25) |
| +7K+12D | 137 | $2.92 \pm 0.05$ | 298 | 0.15 | 7.0 | (25) |
| +7K+12D blocky | 137 | $2.56 \pm 0.05$ | 298 | 0.15 | 7.0 | (25) |
| -4D | 137 | $2.64 \pm 0.05$ | 298 | 0.15 | 7.0 | (25) |
| -8F+4Y | 137 | $2.71 \pm 0.05$ | 298 | 0.15 | 7.0 | (25) |
| -10F+7R+12D | 137 | $2.86 \pm 0.05$ | 298 | 0.15 | 7.0 | (25) |
| +7F-7Y | 137 | $2.72 \pm 0.05$ | 298 | 0.15 | 7.0 | (25) |
| -12F+12Y | 137 | $2.60 \pm 0.05$ | 298 | 0.15 | 7.0 | (25) |
| -12F+12Y-10R | 137 | $2.61 \pm 0.05$ | 298 | 0.15 | 7.0 | (25) |
| -9F+6Y | 137 | $2.65 \pm 0.05$ | 298 | 0.15 | 7.0 | (25) |
| $\alpha$Syn | 140 | $3.55 \pm 0.1$ | 293 | 0.2 | 7.4 | (26) |
| FhuA | 144 | $3.34 \pm 0.1$ | 298 | 0.15 | 7.5 | (24) |
| K17 | 145 | $3.60 \pm 0.2$ | 288 | 0.15 | 7.4 | (27) |
| K27 | 167 | $3.70 \pm 0.2$ | 288 | 0.15 | 7.4 | (27) |
| K10 | 168 | $4.00 \pm 0.1$ | 288 | 0.15 | 7.4 | (27) |
| K16 | 176 | $3.90 \pm 0.3$ | 288 | 0.15 | 7.4 | (27) |
| K25 | 185 | $4.10 \pm 0.2$ | 288 | 0.15 | 7.4 | (27) |
| K32 | 198 | $4.20 \pm 0.3$ | 288 | 0.15 | 7.4 | (27) |
| OPN | 220 | $5.13 \pm 0.2$ | 298 | 0.15 | 6.5 | (28) |
| K23 | 254 | $4.90 \pm 0.2$ | 288 | 0.15 | 7.4 | (27) |
| K44 | 283 | $5.20 \pm 0.2$ | 288 | 0.15 | 7.4 | (27) |
| PNt | 334 | $5.11 \pm 0.2$ | 298 | 0.15 | 7.5 | (24) |

**Table S2. Experimental conditions for the intramolecular PRE data included in the training set.**

| Protein | $N$ | $N_{labels}$ | $\omega_I/2\pi$ (MHz) | $T$ (K) | $c_s$ (M) | pH | Ref. |
|---|---|---|---|---|---|---|---|
| Sic1 | 92 | 6 | 500 | 278 | 0.15 | 7.0 | (29) |
| FUS | 163 | 3 | 850 | 298 | 0.15 | 5.5 | (14) |
| FUS12E | 164 | 3 | 850 | 298 | 0.15 | 5.5 | (14) |
| OPN | 220 | 10 | 800 | 298 | 0.15 | 6.5 | (30) |
| aSyn | 140 | 5 | 700 | 283 | 0.15 | 7.4 | (11) |
| A2 | 155 | 2 | 850 | 298 | 0.005 | 5.5 | (15) |

**Table S3. Experimental conditions for the intermolecular PRE data included in the test set.**

| Protein | $N$ | $N_{labels}$ | $\omega_I/2\pi$ (MHz) | $T$ (K) | $c_s$ (M) | pH | [Protein] µM | Ref. |
|---|---|---|---|---|---|---|---|---|
| FUS | 163 | 3 | 850 | 298 | 0.15 | 5.5 | 50 | (14) |
| A2 | 155 | 2 | 850 | 298 | 0.005 | 5.5 | 30 | (15) |

**Table S4. Solution conditions of slab simulations.**

| Protein | $c_s$ (mM) | pH | Ref. | T (K) | | Software |
|---|---|---|---|---|---|---|
| | | | | HPS & M1–3 | HPS-Urry | |
| A2 LCD WT | 10 | 5.5 | (15, 31) | 323 | 297 | HOOMD-blue |
| A2 LCD NtoS | 10 | 5.5 | (31) | 323 | 297 | HOOMD-blue |
| Ddx4 LCD WT and CS | 130 | 6.5 | (32–34) | 323 | 297 | HOOMD-blue & openMM |
| Ddx4 LCD CS, FA and RK | 130 | 6.5 | (32–34) | 297 | 297 | HOOMD-blue & openMM |
| A1 LCD WT & Variants | 150 | 7.0 | (25, 35) | 323 & 310 | 297 | HOOMD-blue |
| FUS LCD | 150 | 5.5 | (36, 37) | 323 | 297 | HOOMD-blue |
| LAF-1 RGG domain | 150 | 7.5 | (38–40) | 323 | 297 | HOOMD-blue & openMM |
| LAF-1 RGG domain | 80 | 7.5 | (38–40) | 323 (M1 only) | – | openMM |
| LAF-1 RGG domain Variants | 150 | 7.5 | (38–40) | 323 (M1 only) | – | openMM |

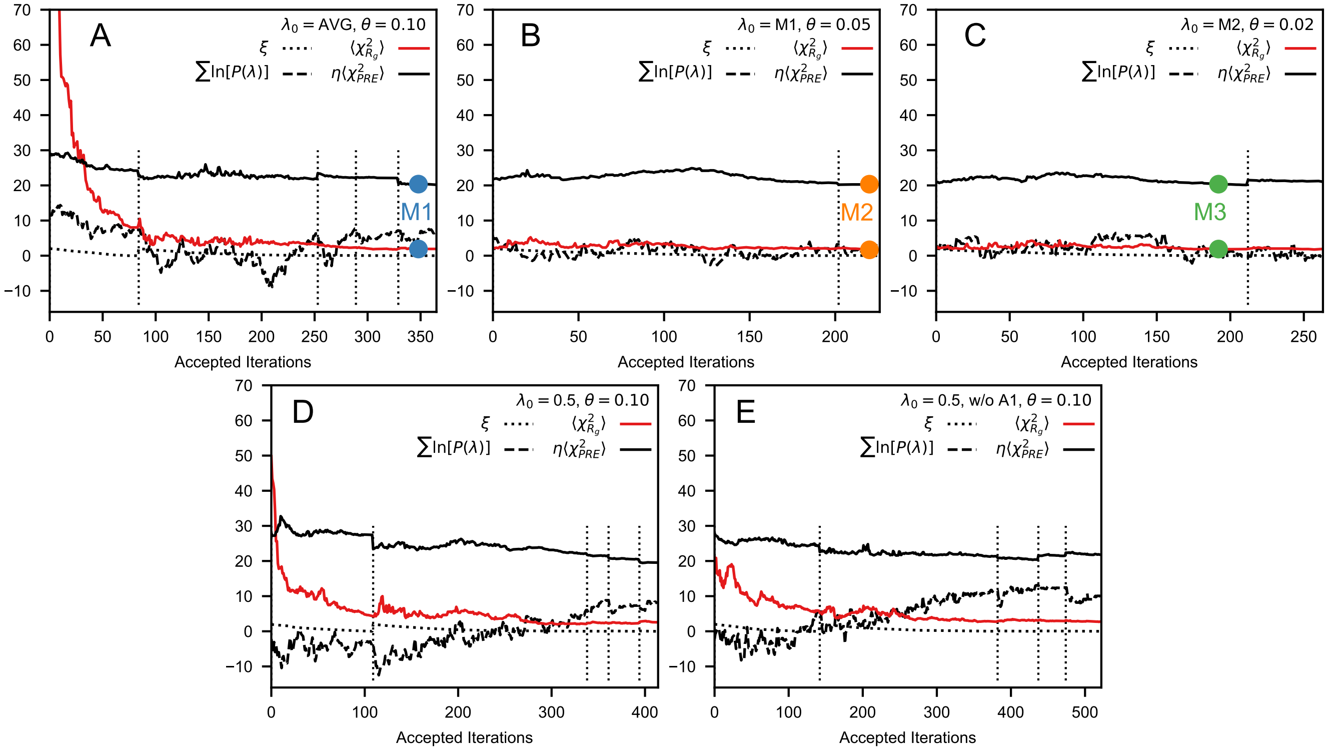


**Fig. S1.** Evolution of the control parameter $\xi$ (black dotted line), the discrepancy between $\lambda$ set and prior $\sum_i \ln\left[P(\lambda_i)\right]$ (black dashed line), $\chi 2_{R_g}$ (red solid) and $\chi 2_{PRE}$ (black solid line) during the optimization procedures performed with $\theta = 0.1$ (*A*, *D* and *E*), 0.05 (*B*) and 0.02 (C), starting from $\lambda_0$ =AVG (*A*), M1 (*B*), M2 (*C*) and 0.5 (*D* and *E*). Dotted vertical lines indicate updated sampling by molecular simulations, whereas the remaining points are estimated from the reweighted ensembles.

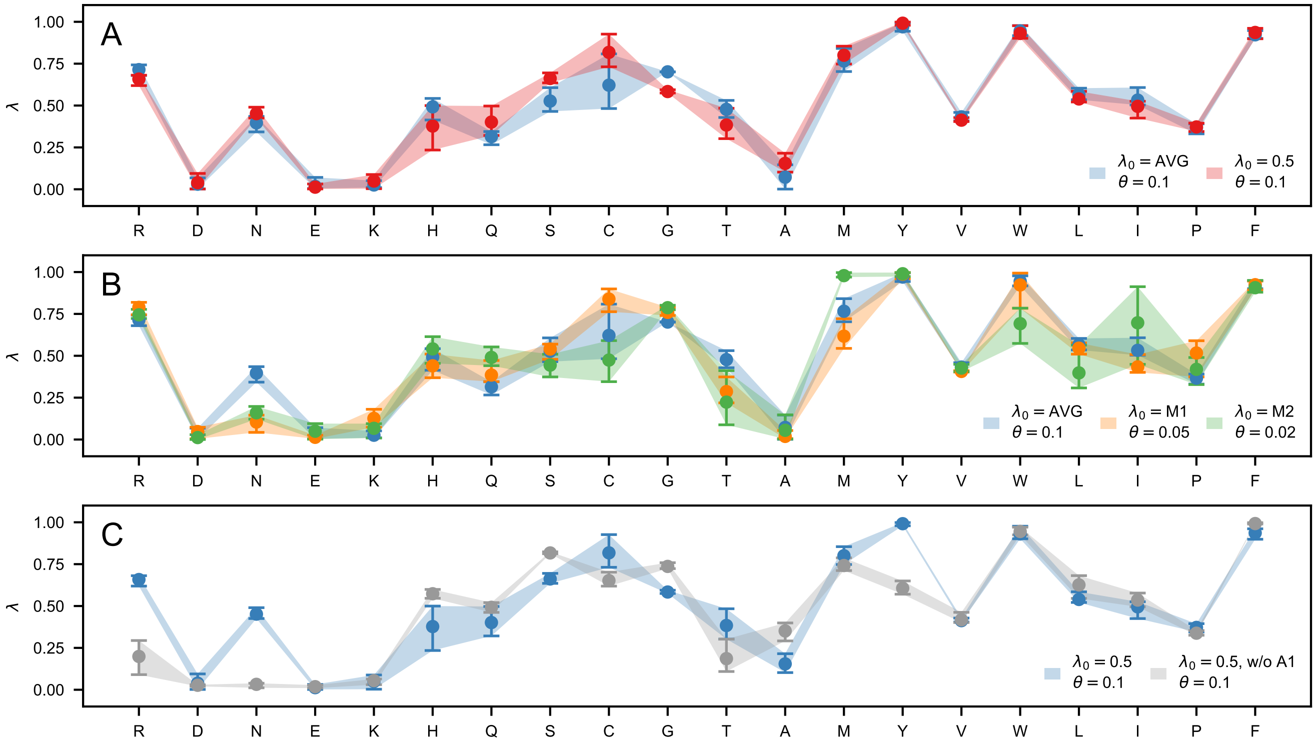


**Fig. S2.** Comparison between ranges of optimal $\lambda$ values collected through the optimization procedures performed in this study. Circles show mean values whereas error bars and shaded areas show the interval between 10% and 90% quantiles. (*A*) Optimization procedures performed using the same hyperparameters result in largely overlapping ranges of optimal $\lambda$ values, irrespective of whether the starting point is the AVG values (blue) or $\lambda = 0.5$ (red). (*B*) Decreasing the confidence parameter results in an overall expansion of the range of the optimal $\lambda$ values. (*C*) We observe significant differences between optimal $\lambda$ values obtained using the whole training set and excluding the 21 variants of A1 LCD. Notably, the A1 LCD variants are responsible for the large hydrophobicity estimated for Y and R.

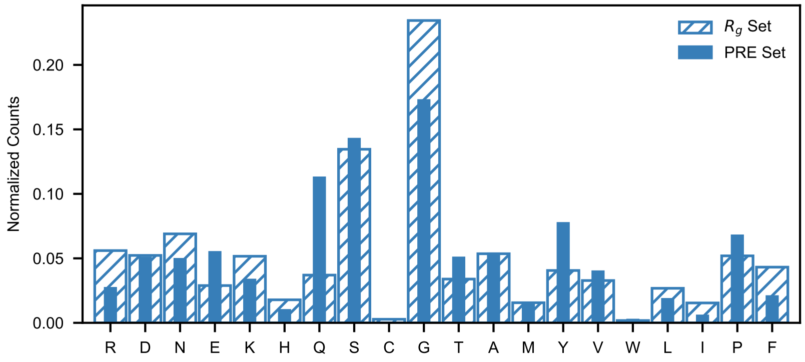


**Fig. S3.** Amino acid probability distributions for proteins in the $R_g$ (hatched) and in the PRE training sets (closed).

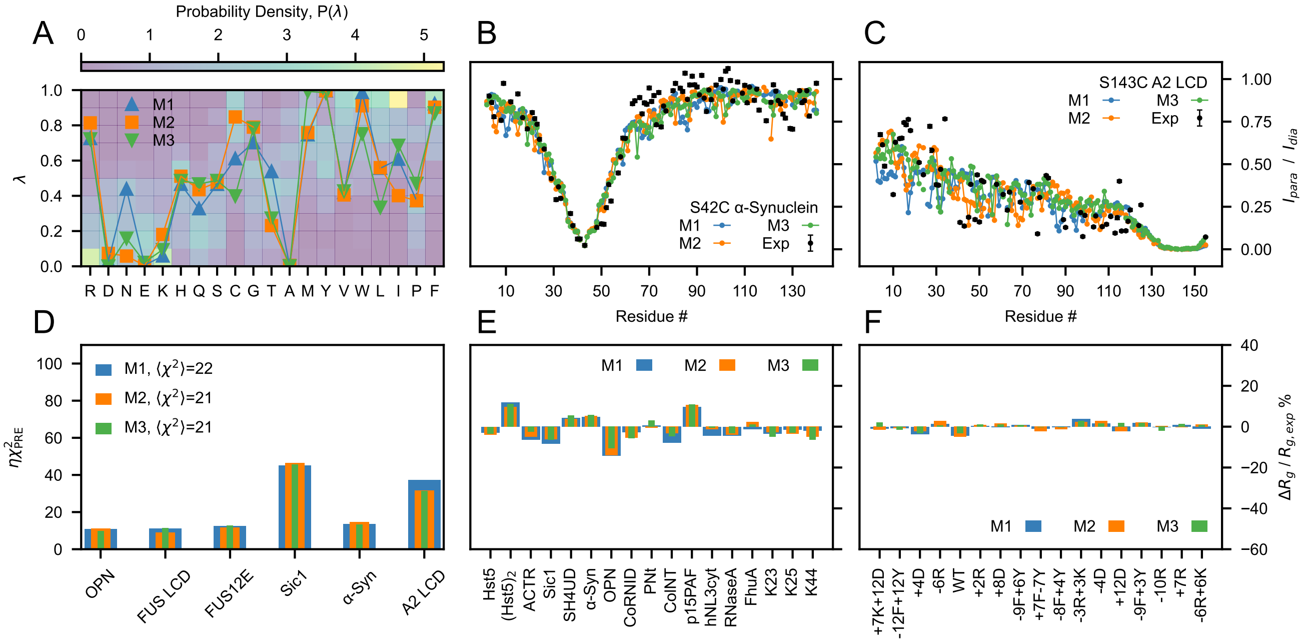


**Fig. S4.** (*A*) Probability distributions of the $\lambda$ parameters calculated from 87 min-max normalized hydrophobicity scales. The overlaying lines are the $\lambda$ parameters of the M1 (blue), M2 (orange) and M3 (green) models. Intramolecular PRE rates for the S43C mutant of $\alpha$-Synuclein (*B*) and the S243C mutant of A2 LCD (*C*) from simulations of models M1–3 and from experiments (11, 15) (black). (*D*) $\chi^2$ values quantifying the discrepancy between simulated and experimental intermolecular PRE data. Relative difference between simulated and experimental gyration radii for non phase-separating sequences (*E*) and for variants of A1 LCD (*F*).

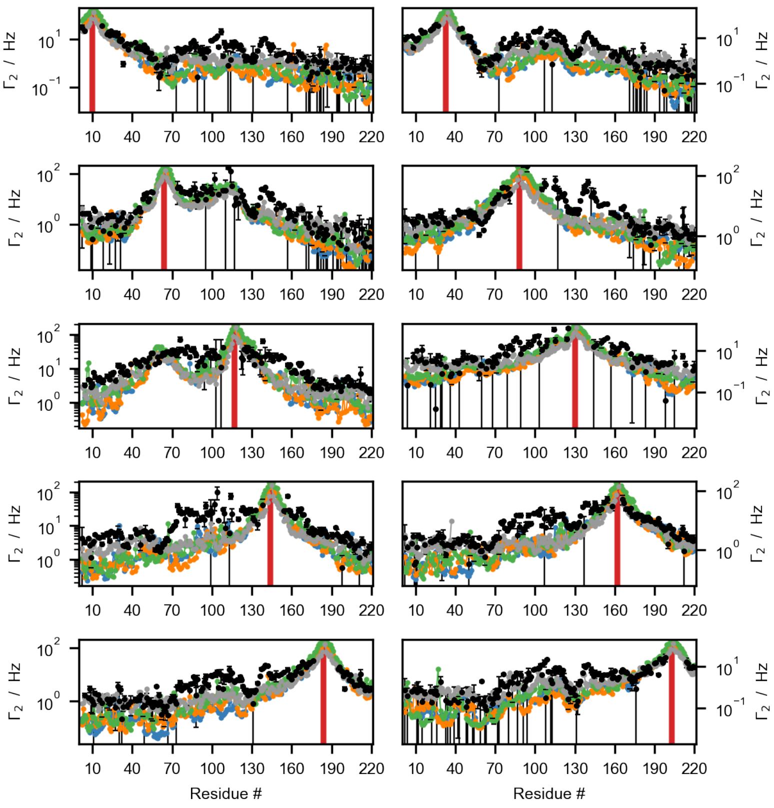


**Fig. S5.** Comparison between experimental PRE rates (black) for the S10C, E33C, S64C, R88C, A117C, D130C, S144C, S162C, S184C and S203C mutants of OPN (30) and corresponding values calculated from the M1 (blue), M2 (orange), M3 (green) and HPS (gray) models using $\tau_c = 2$, 2, 3 and 1 ns, respectively. Red vertical lines indicate spin-labeled positions.

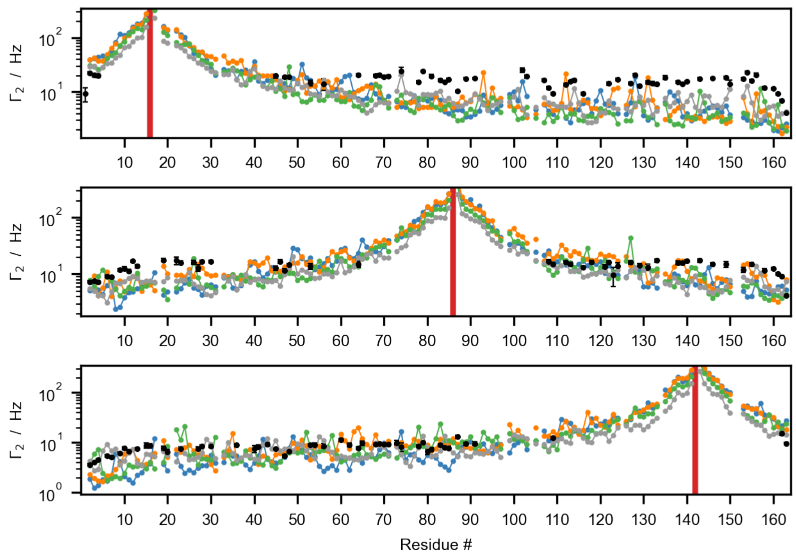


**Fig. S6.** Comparison between experimental PRE rates (black) for the A16C, S86C and S142C mutants of FUS LCD (14) and corresponding values calculated from the M1 (blue), M2 (orange), M3 (green) and HPS (gray) models using $\tau_c = 4$, 4, 3 and 2 ns, respectively. Red vertical lines indicate spin-labeled positions.

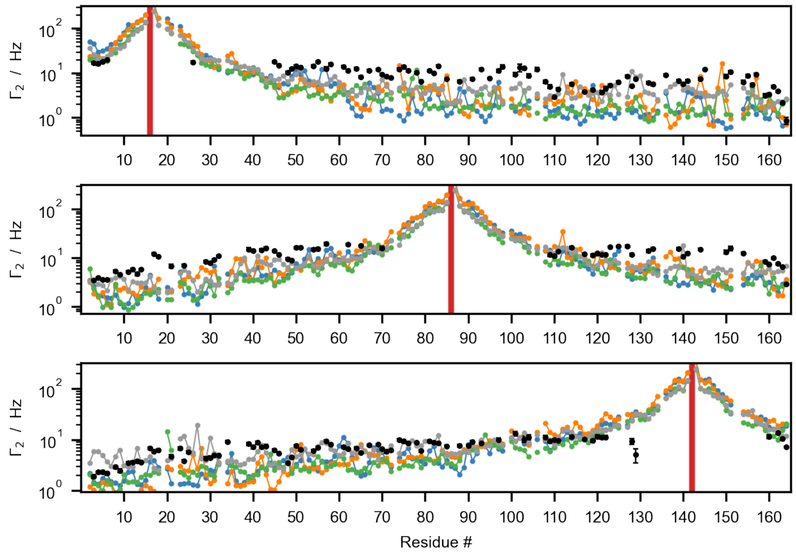


**Fig. S7.** Comparison between experimental PRE rates (black) for the A16C, S86C and S142C mutants of FUS12E LCD (14) and corresponding values calculated from the M1 (blue), M2 (orange), M3 (green) and HPS (gray) models using $\tau_c = 3$, 3, 2 and 2 ns, respectively. Red vertical lines indicate spin-labeled positions.

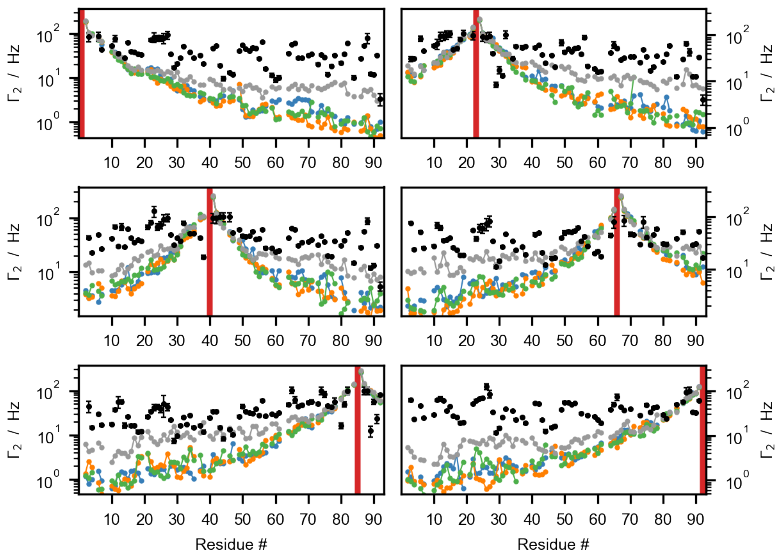


**Fig. S8.** Comparison between experimental PRE rates (black) for the G1C, N23C, S40C, N66C, P85C and T92C mutants of Sic1 (29) and corresponding values calculated from the M1 (blue), M2 (orange), M3 (green) and HPS (gray) models using $\tau_c = 2$ ns. Red vertical lines indicate spin-labeled positions.

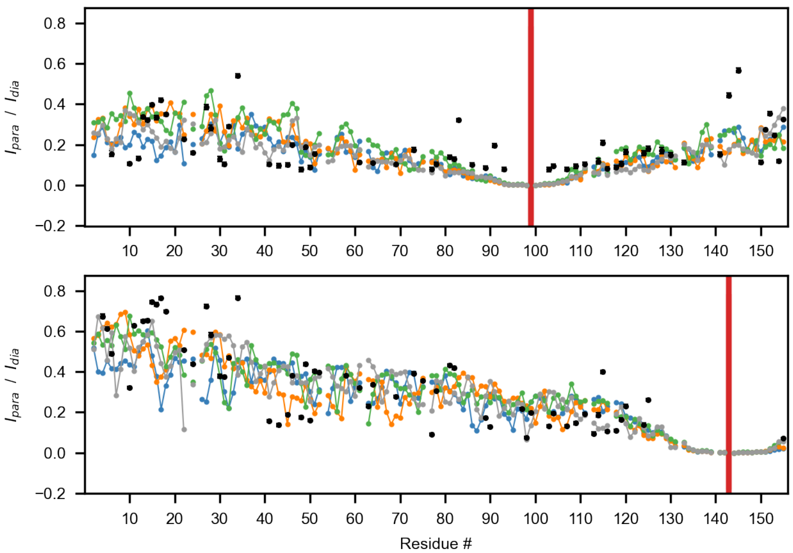


**Fig. S9.** Comparison between experimental PRE ratios (black) for the S99C and S143C mutants of A2 LCD (15) and corresponding values calculated from the M1 (blue), M2 (orange), M3 (green) and HPS (gray) models using $\tau_c = 6$, 6, 5 and 6 ns, respectively. Red vertical lines indicate spin-labeled positions.

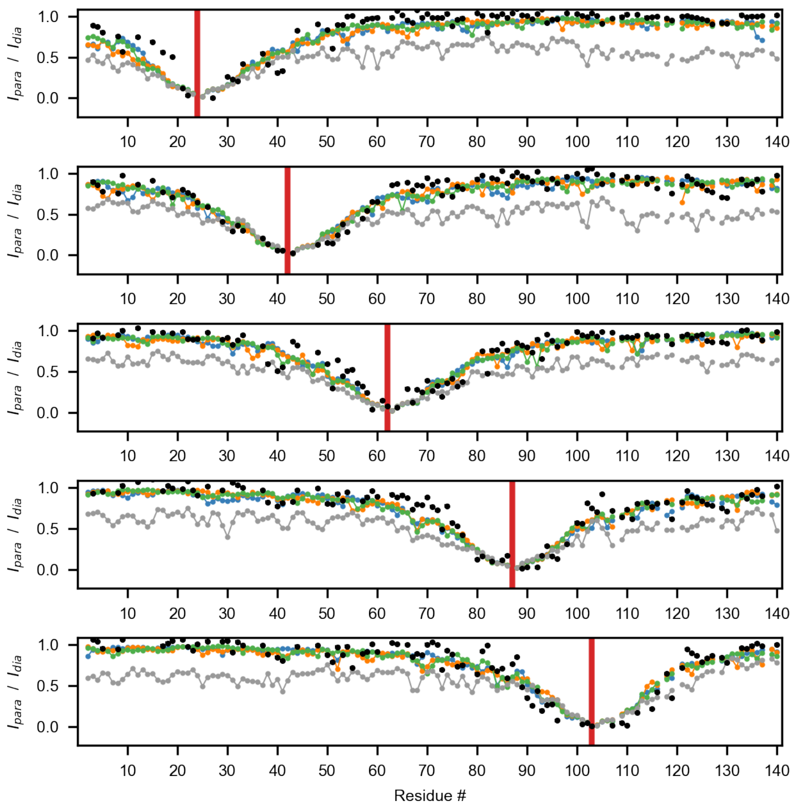


**Fig. S10.** Comparison between experimental PRE ratios (black) for the Q24C, S42C, Q62C, S87C and N103C mutants of $\alpha$-Synuclein (11) and corresponding values calculated from the M1 (blue), M2 (orange), M3 (green) and HPS (gray) models using $\tau_c = 1$ ns. Red vertical lines indicate spin-labeled positions.

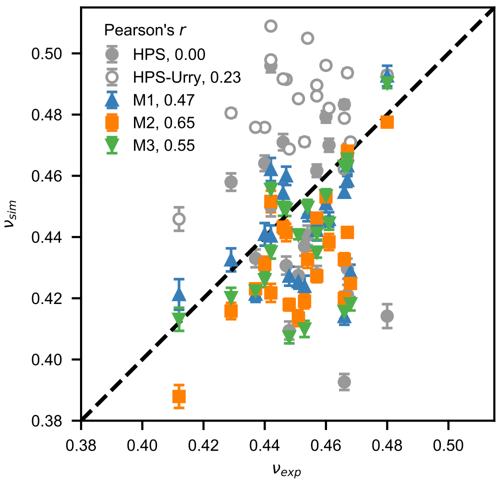


**Fig. S11.** Correlation between experimental and predicted scaling exponents, $\nu$, for the A1 LCD variants in Tab. S1 simulated using the HPS, HPS-Urry and M1–3 models.

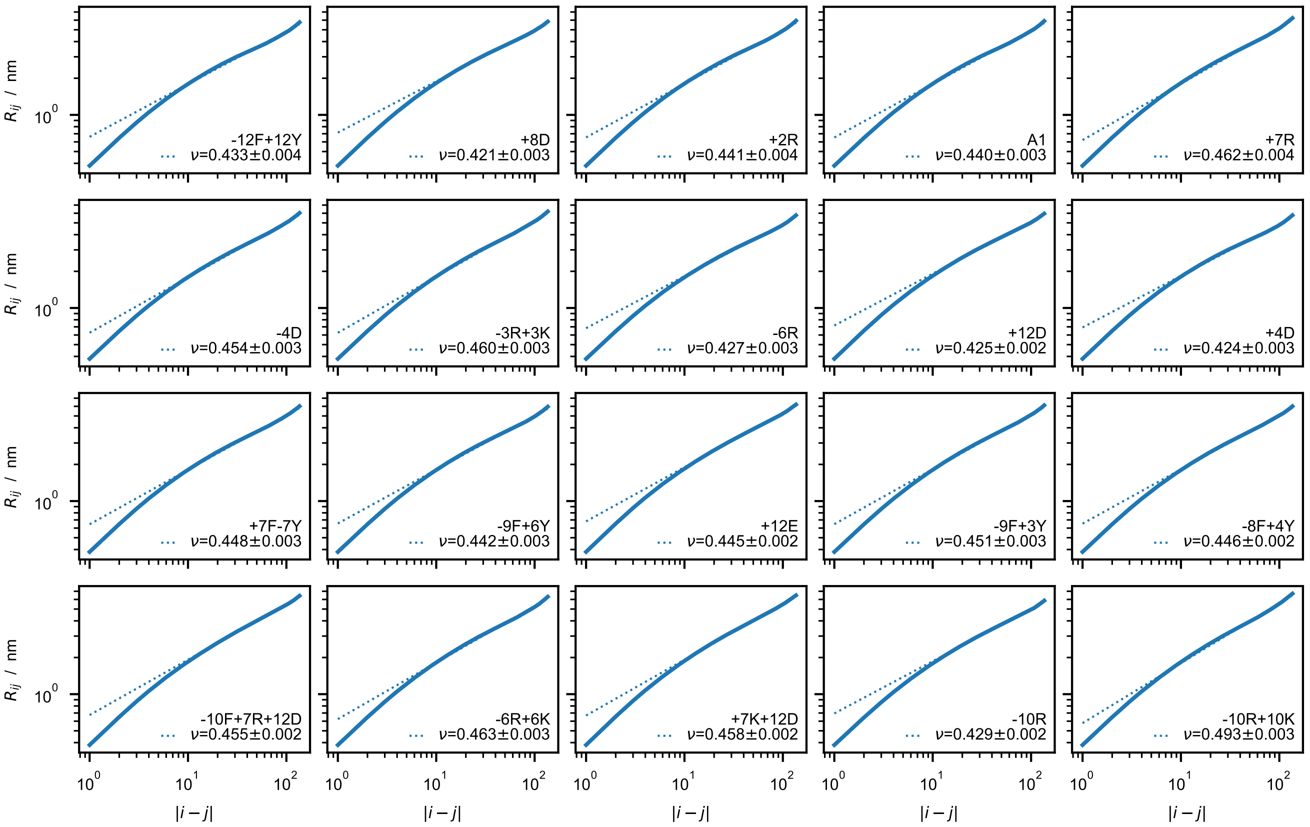


**Fig. S12.** Calculation of the scaling exponent, $\nu$, from single-chain simulations with the M1 model at 25°C. Log-log plots of the intramolecular pairwise distances, $R_{ij}$, as a function of the absolute distance of the residues along the chain $|i-j|$ (solid lines). The long-distance region, $|i-j| > 10$, is fitted to $R_{ij} = R_0|i-j|^{\nu_{sim}}$ (dotted lines) where $R_0$ is a constant and $\nu_{sim}$ is the apparent scaling exponent.

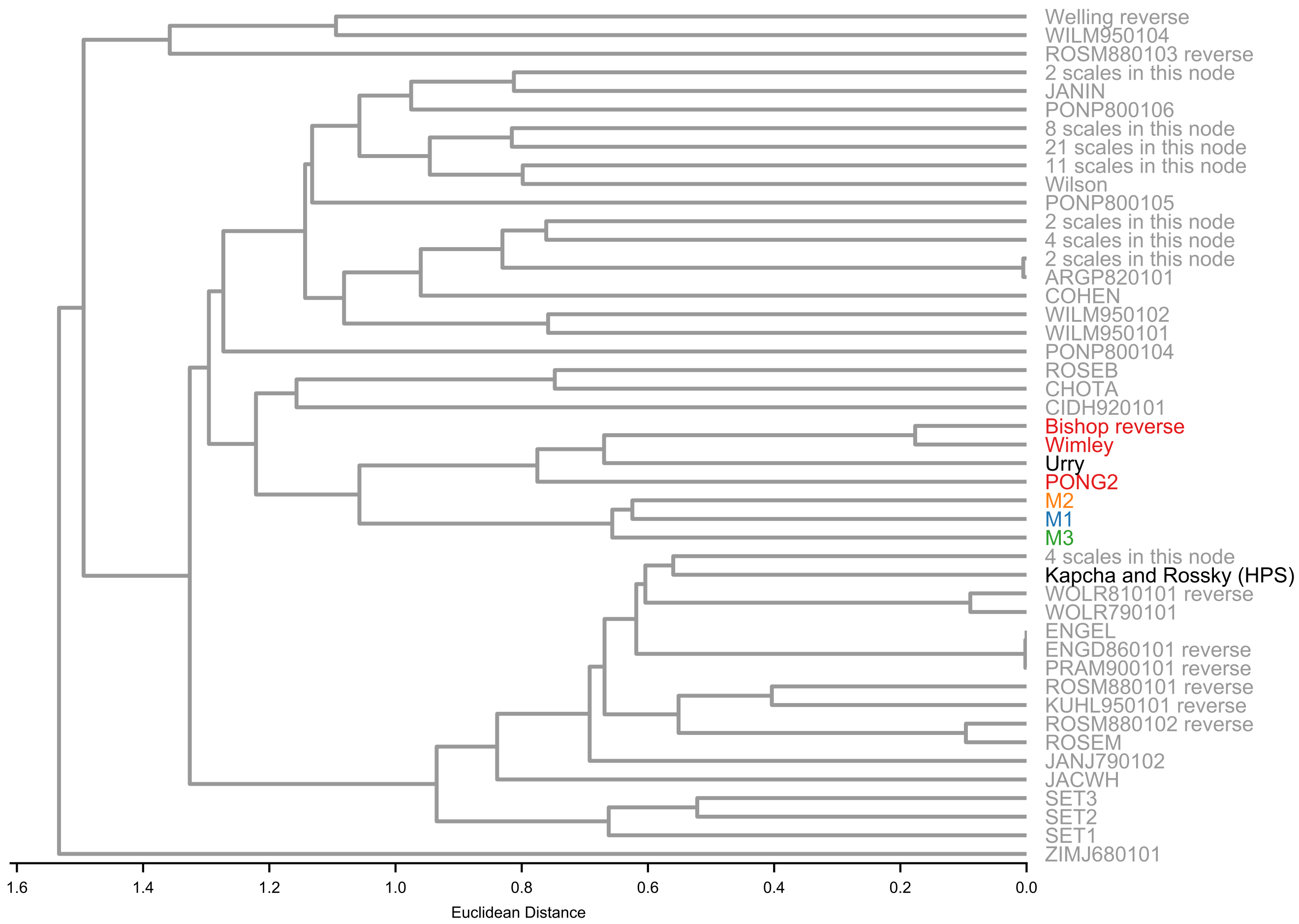


**Fig. S13.** Hierarchical clustering dendrogram of min-max normalized hydrophobicity scales including the 87 scales retained from the set by Simm et al. (41), the M1–3 scales proposed in this work (blue, orange and green labels) as well as the HPS and Urry scales (black labels). Red labels highlight the scales with closest linkage distance from M1–3. The agglomerative clustering is performed using Euclidean distances and the average linkage method as implemented in the Python scikit-learn package (42).

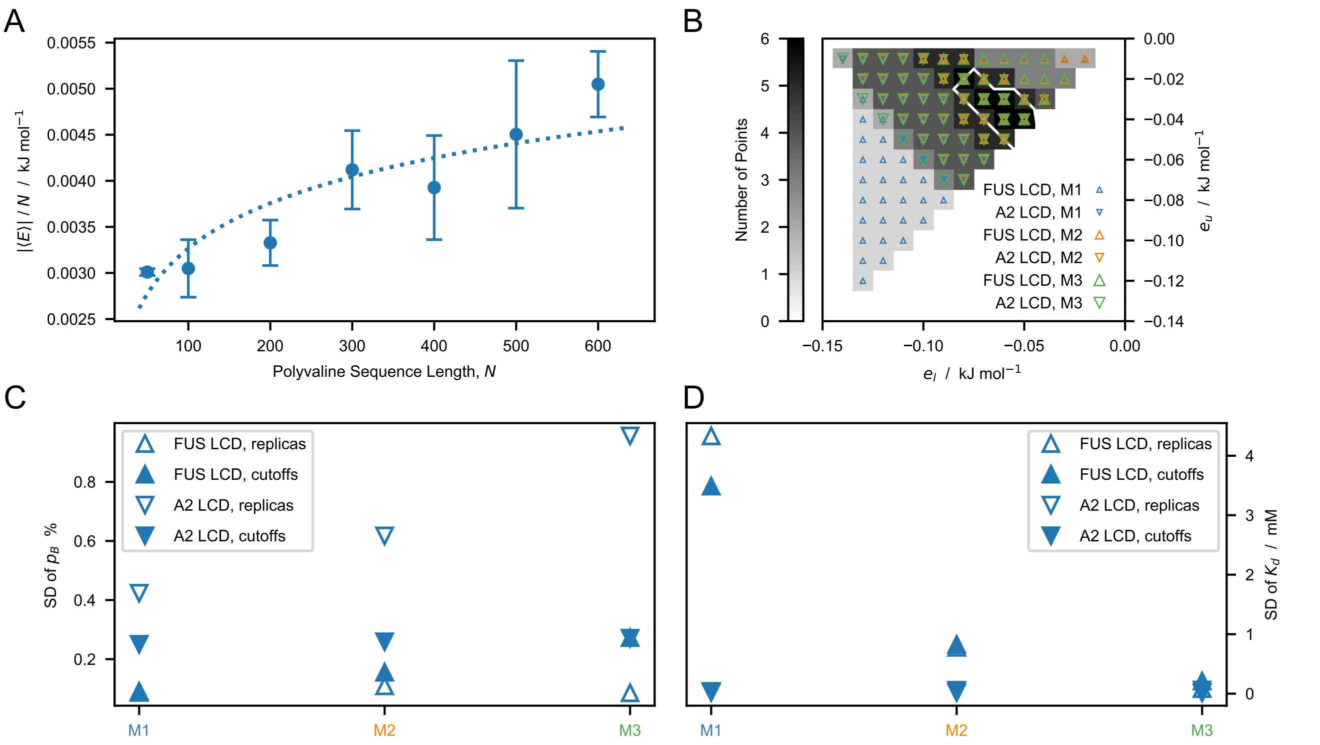


**Fig. S14.** (*A*) $N$-normalized average intermolecular non-electrostatic interaction energy between polyvaline chains of various sequence length, $N$. Error bars are SDs of three simulation replicas of 2 µs each. The dotted line is the non-linear least-squares fit to $c \times \ln N$ where c is a constant. (*B*) Dual cutoffs, $[e_l,\, e_u]$, resulting in $2(K_d - K_d')/(K_d + K_d') < 5\%$ for FUS LCD (△) and A2 LCD (▽) modeled using M1 (blue), M2 (orange) and M3 (green). The color gradient shows the overall distribution of cutoff values whereas the white contour line encloses the region of maximal overlap used to identify the optimal dual cutoffs. Comparison between SDs of $p_B$ (*C*) and $K_d$ (*D*) estimated from ten simulation replicas with $[e_l, e_u]$=[-0.07,-0.05] kJ mol$^{-1}$ (open symbols) or from the five optimal cutoffs [-0.09,-0.03], [-0.08,-0.04], [-0.07,-0.04], [-0.07,-0.05] and [-0.06,-0.05] kJ mol$^{-1}$ (closed symbols).

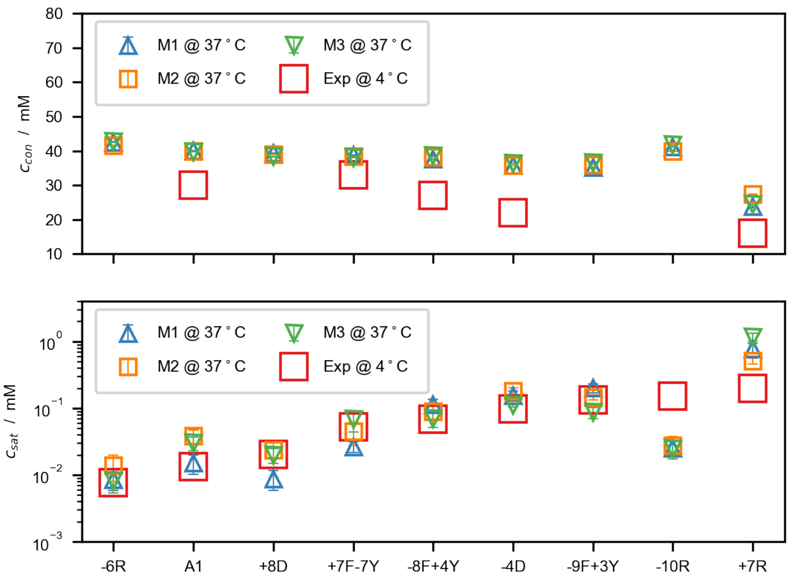


**Fig. S15.** Protein concentrations in the condensate (*Top*) and in the dilute phase (*bottom*) from slab simulations of the M1–3 models at 37°C. Red open squares indicate experimental measurements at ∼4°C (25). Error bars are SEMs of averages over blocks of 0.3 µs.

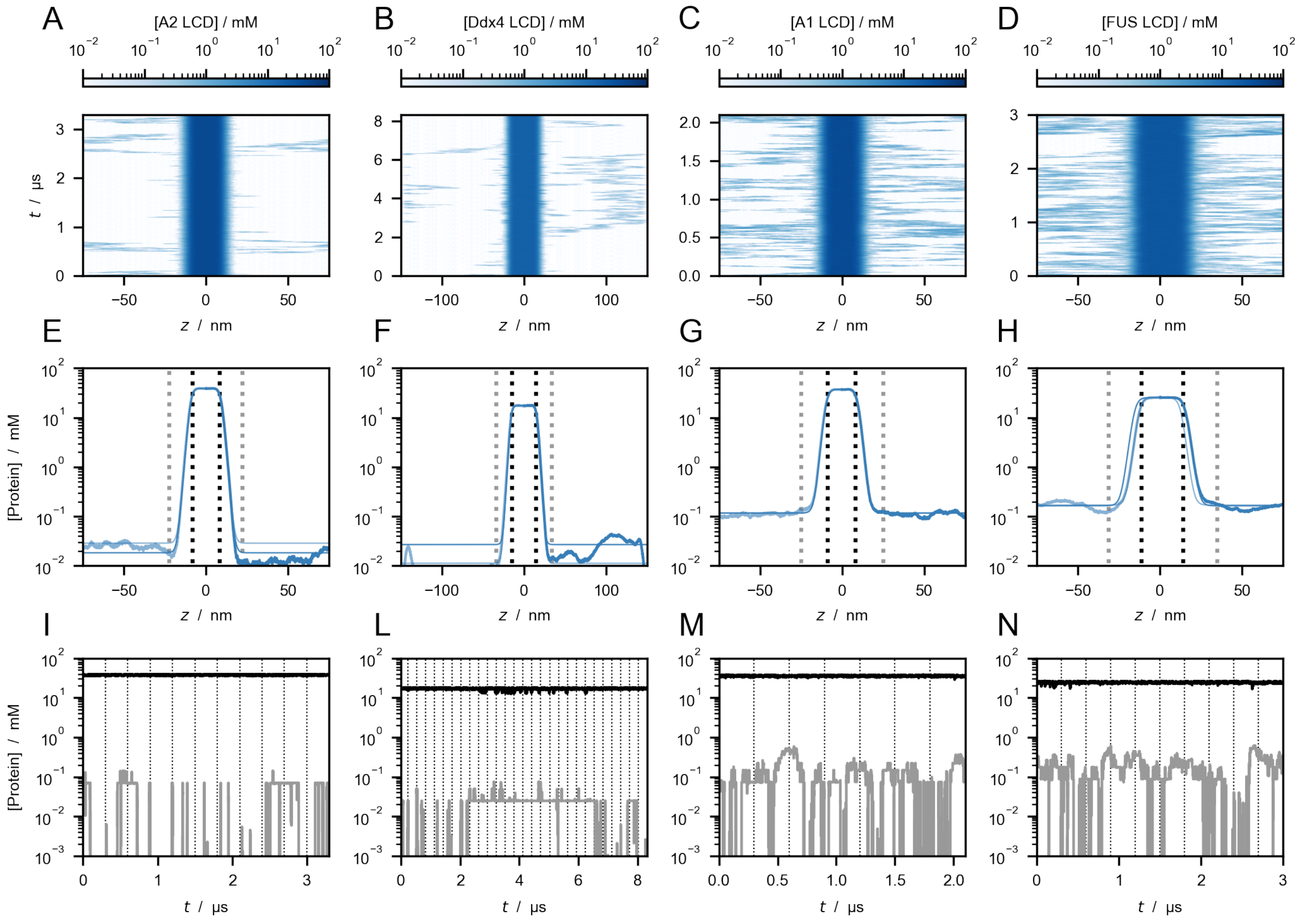


**Fig. S16.** (*A*–*D*) Time evolution of the protein concentration profiles from slab simulations of the M1 model at 50°C of A2 LCD (*A*), Ddx4 LCD (*B*), A1 LCD (*C*) and FUS LCD (*D*). (*E*–*H*) Concentration profiles. (*I*–*N*) Time series of the protein concentration in the slab (black) and in the dilute phase (gray).

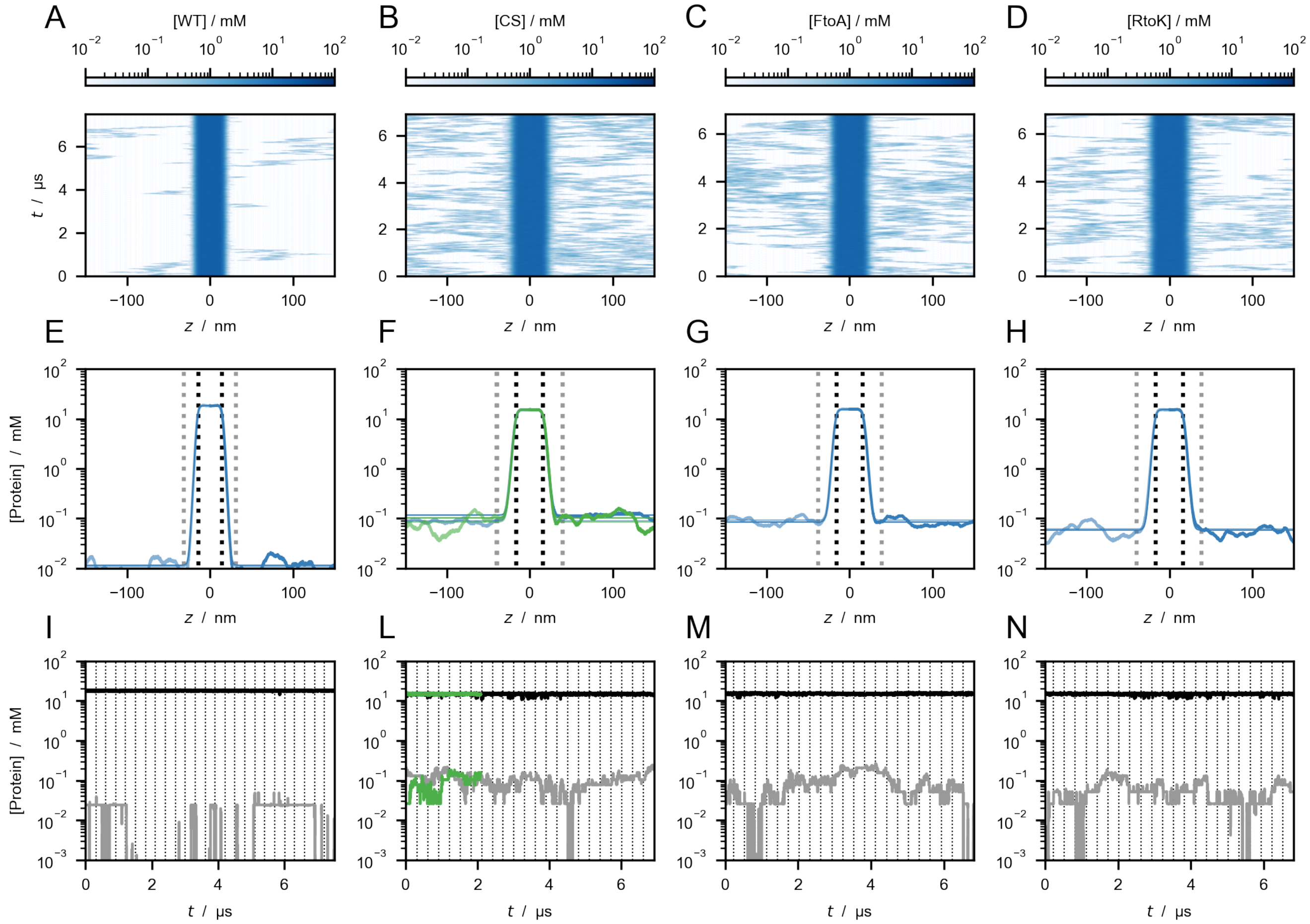


**Fig. S17.** (*A*–*D*) Time evolution of the protein concentration profiles from slab simulations of the HPS-Urry model at 24°C of Ddx4 LCD WT (*A*), CS (*B*), FtoA (*C*) and RtoK (*D*). (*E*–*H*) Concentration profiles. (*I*–*N*) Time series of the protein concentration in the slab (black) and in the dilute phase (gray). Green lines in *L* are obtained from simulations performed using HOOMD-blue whereas the remainder data shown in this figure are obtained using openMM.

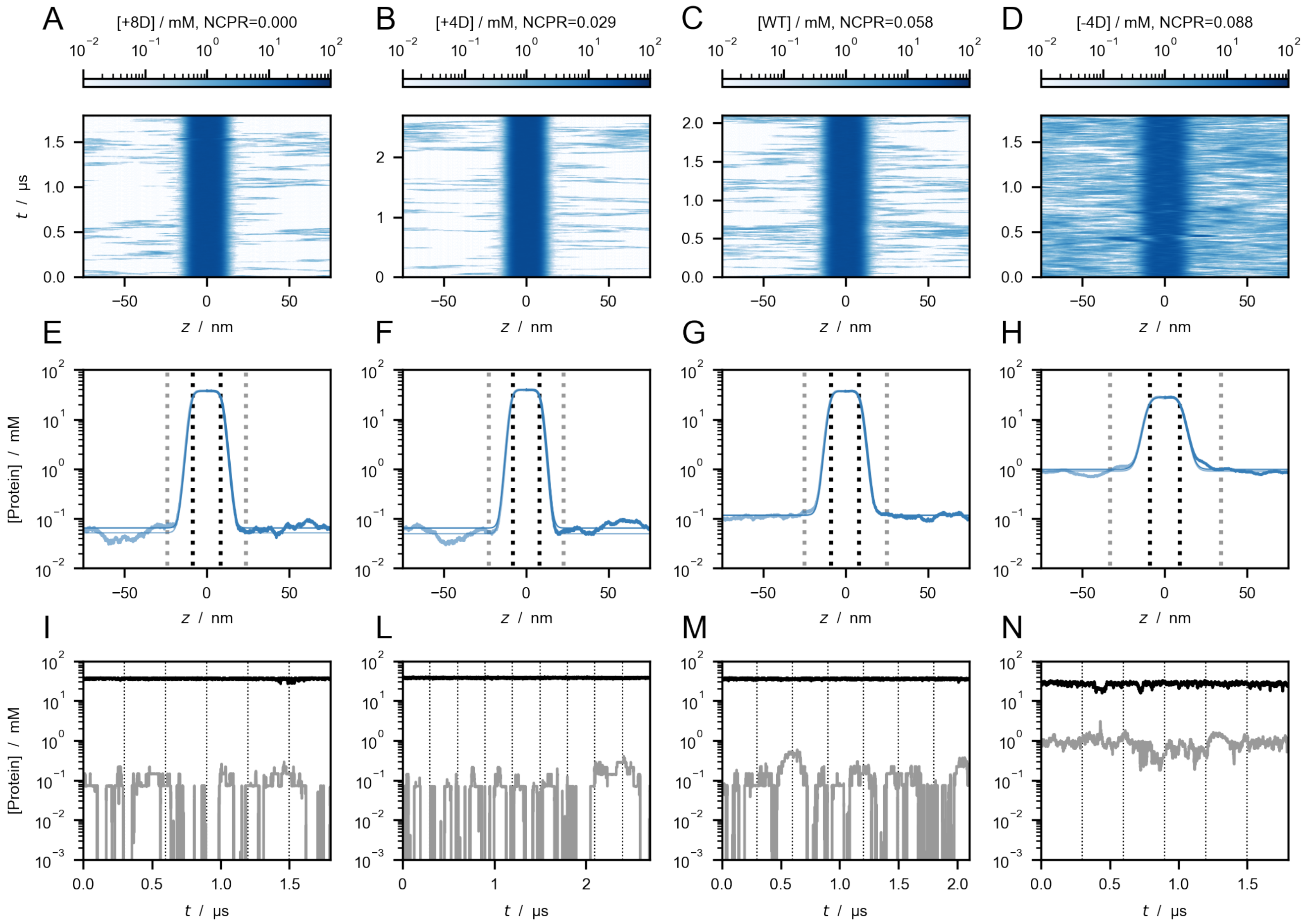


**Fig. S18.** (*A–D*) Time evolution of the protein concentration profiles from slab simulations of the M1 model at 50°C of the +8D (*A*), +4D (*B*), WT (*C*) and -4D (*D*) variants of A1 LCD. (*E–H*) Concentration profiles. (*I–N*) Time series of the protein concentration in the slab (black) and in the dilute phase (gray).

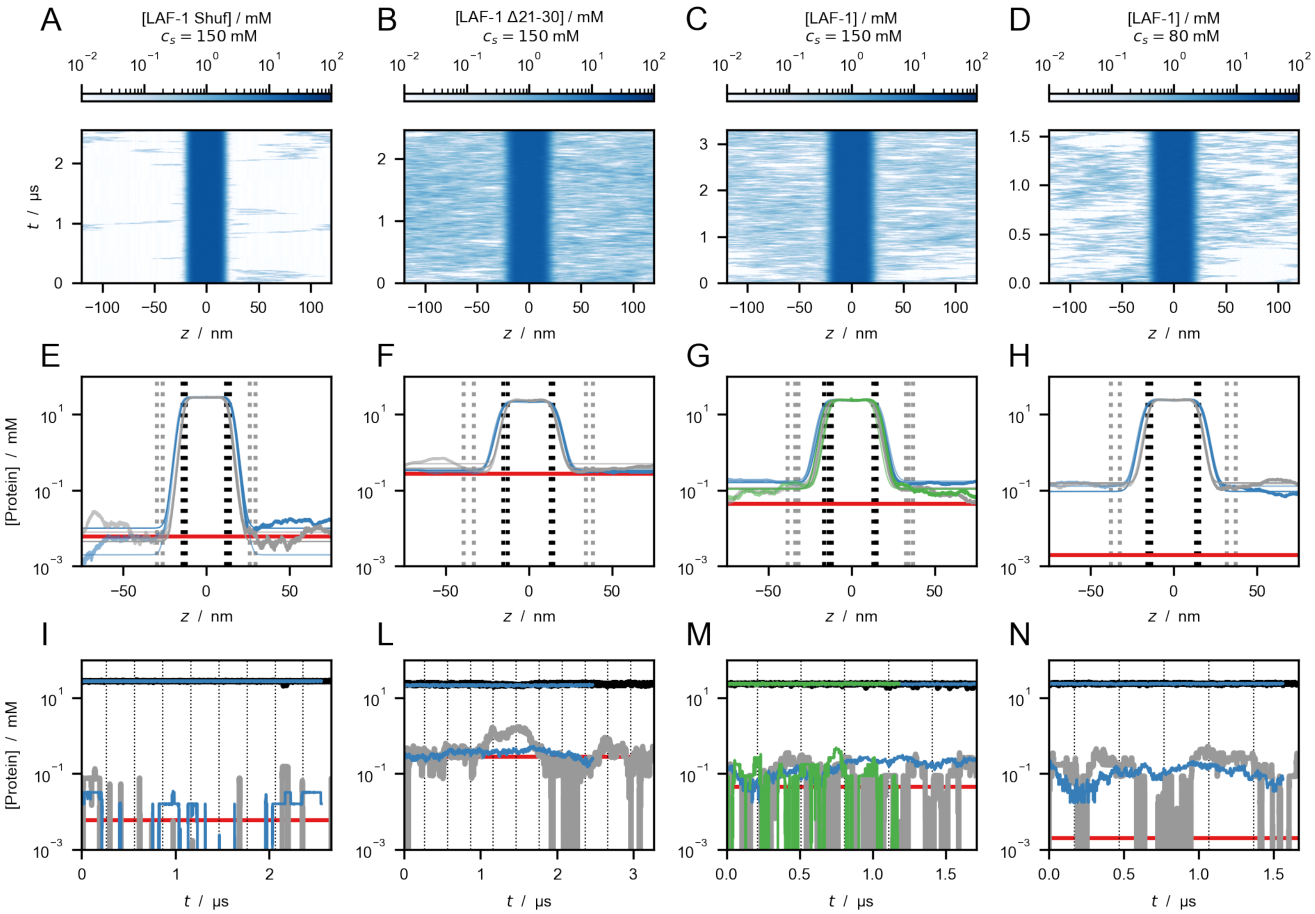


**Fig. S19.** (*A*–*D*) Time evolution of the protein concentration profiles from 300-chain slab simulations of the M1 model at 50°C of the shuffled (*A*), Δ21–30 (*B*), and WT (*C* and *D*) variants of LAF-1 RGG domain. All the simulations are performed at $c_s = 150$ mM and the WT is further simulated at $c_s = 80$ mM (*D*). (*E*–*H*) Concentration profiles from 100-chain (gray) and 300-chain (blue) slab simulations. (*I*–*N*) Time series of the protein concentration in the slab (black) and in the dilute phase (gray) from 100-chain simulations. Blue lines show the corresponding time series from 300-chain simulations. Red horizontal lines indicate the experimental $c_{sat}$ values reported by Schuster et al. (40) (*E*, *F*, *G*, *I*, *L* and *M*) and Taylor et al. (39) (*H* and *N*). Green lines in *G* and *M* are obtained from simulations performed using HOOMD-blue whereas the remainder data shown in this figure are obtained using openMM.

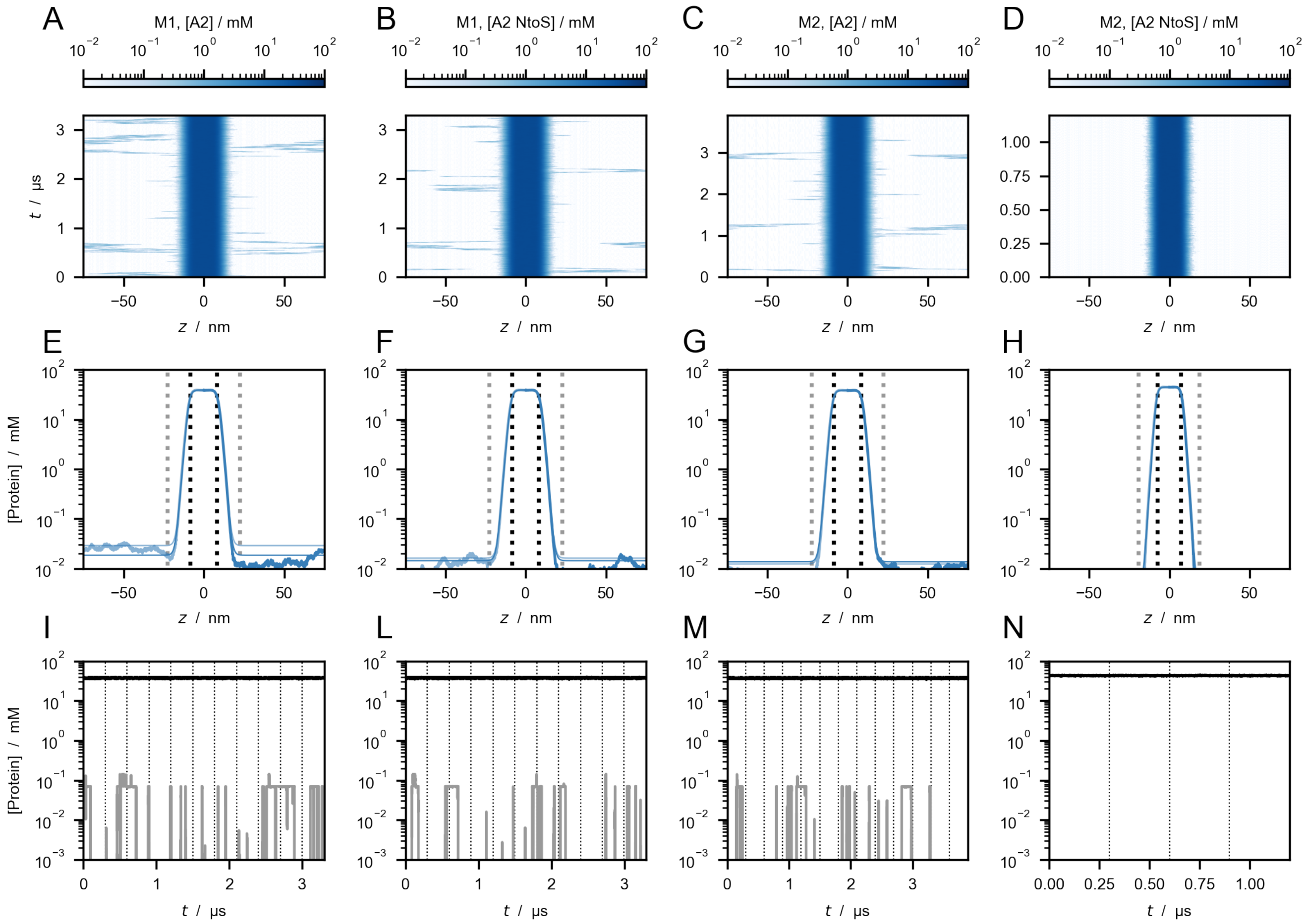


**Fig. S20.** (*A*–*D*) Comparison between the M1 and M2 models for the time evolution of the protein concentration profiles from slab simulations of A2 LCD WT (*A* and *C*) and NtoS vairant (*B* and *D*). (*E*–*H*) Concentration profiles. (*I*–*N*) Time series of the protein concentration in the slab (black) and in the dilute phase (gray).

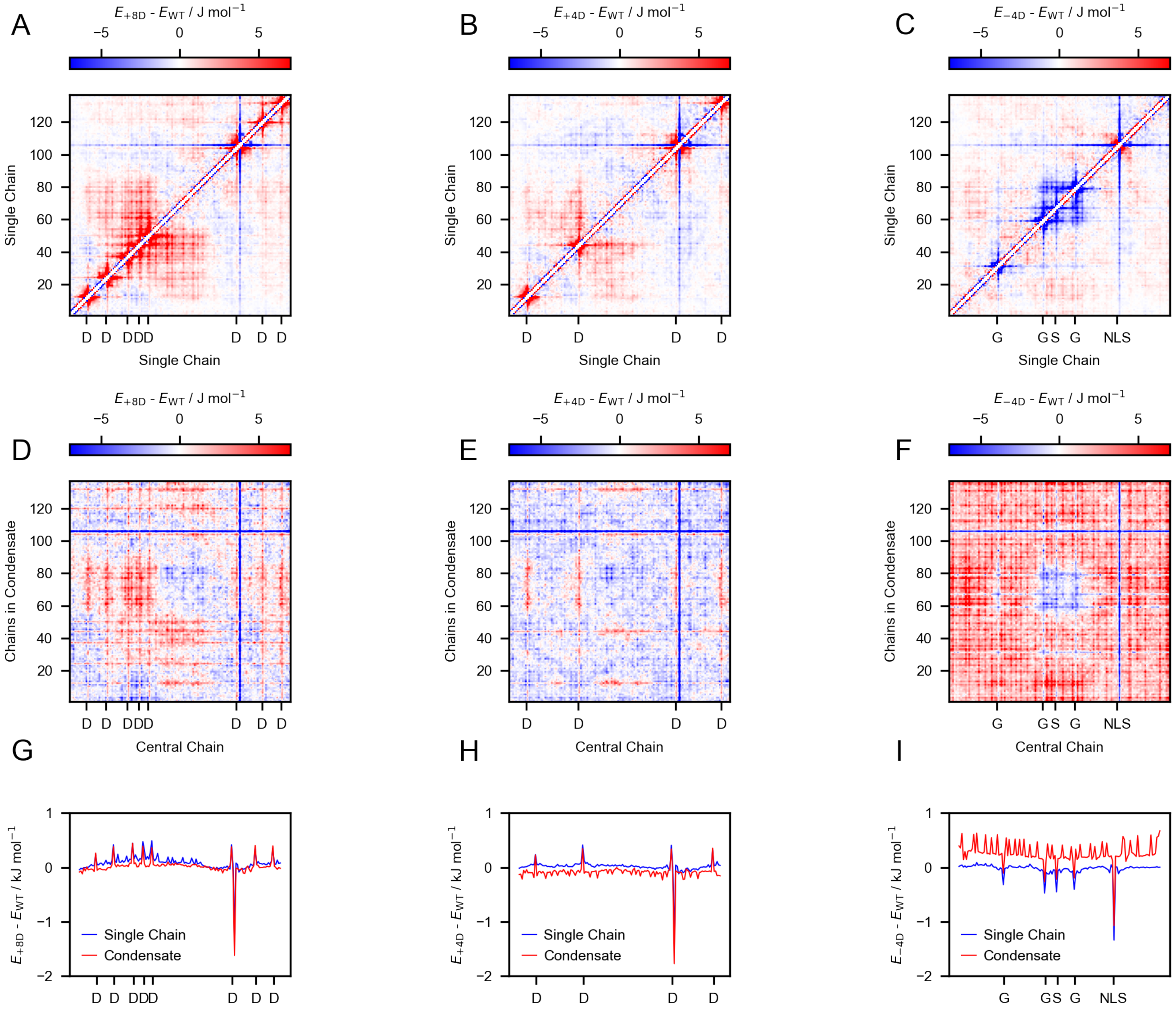


**Fig. S21.** Residual energy maps of +8D, +4D and -4D variants with respect to the wild type of A1 LCD. (*A*–*C*) Residual energy maps from single-chain simulations of the M1 model at 25°C. (*D*–*F*) Residual energy maps from slab simulations with the M1 model at 50 °C where the non-electrostatic pairwise interactions energies are calculated between a chain located at the center of a condensate and the surrounding chains. (*G*–*I*) 1D projections of the residual energy maps calculated within a single chain (blue) and among distinct chains in the condensate (red) for +8D (NCPR=0), +4D (NCPR≈0.03) and -4D (NCPR≈0.09) variants with respect to WT (NCPR≈0.06). The large energy differences for residues 104 and 105 are due to the PY nuclear localization signal (PY-NLS) which is present in the variants whereas it is replaced by GS in the WT.

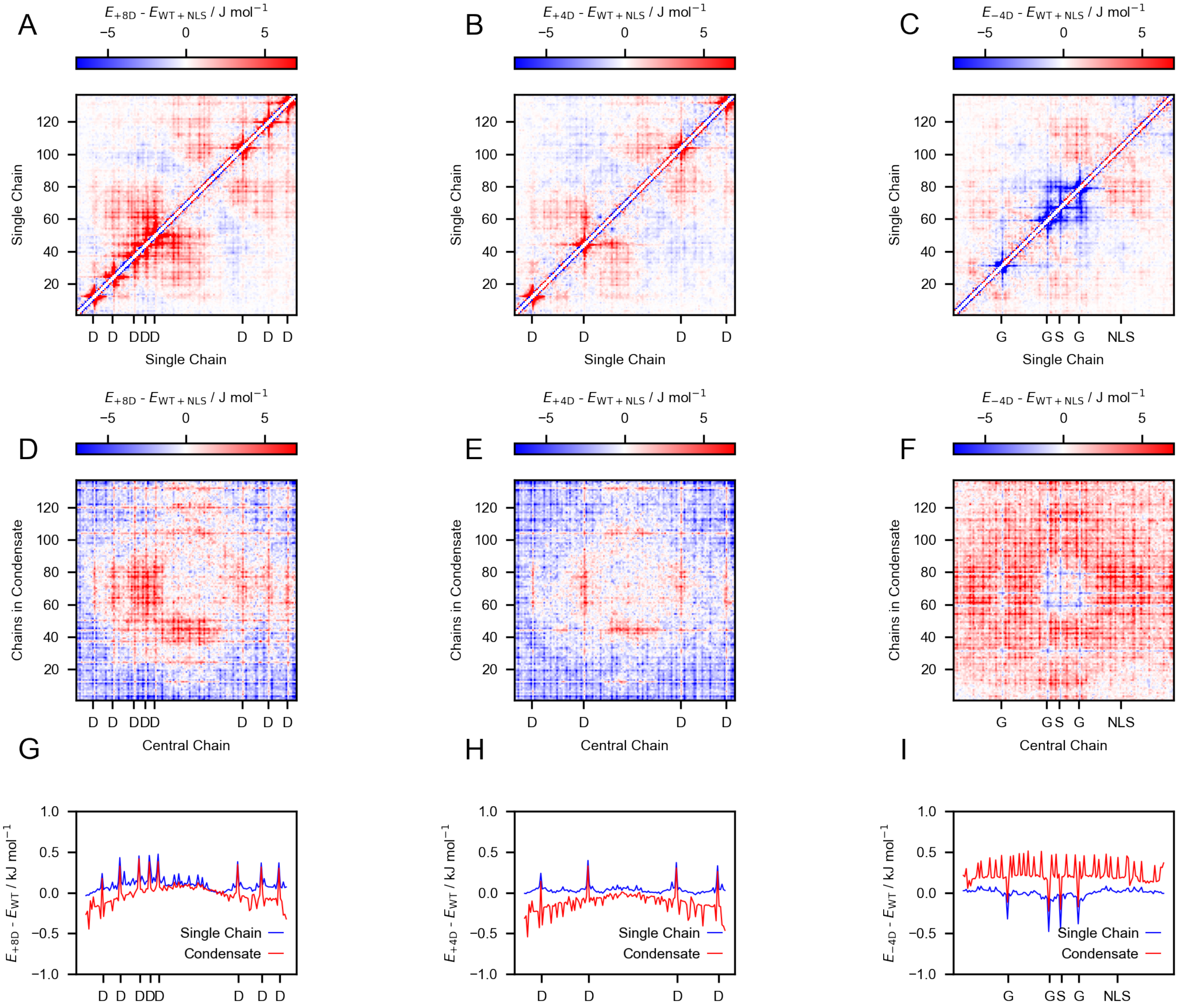

**Fig. S22.** Residual energy maps of +8D, +4D and -4D variants with respect to the wild type of A1 LCD with the PY nuclear localization signal (WT+NLS). (*A–C*) Residual energy maps from single-chain simulations with the M1 model at 25°C. (*D–F*) Residual energy maps from slab simulations with the M1 model at 50°C where the non-electrostatic pairwise interactions energies are calculated between a chain located at the center of a condensate and the surrounding chains. (*G–I*) 1D projections of the residual energy maps calculated within a single chain (blue) and among distinct chains in the condensate (red) for +8D (NCPR=0), +4D (NCPR≈0.03) and -4D (NCPR≈0.09) variants with respect to WT+NLS (NCPR≈0.06).

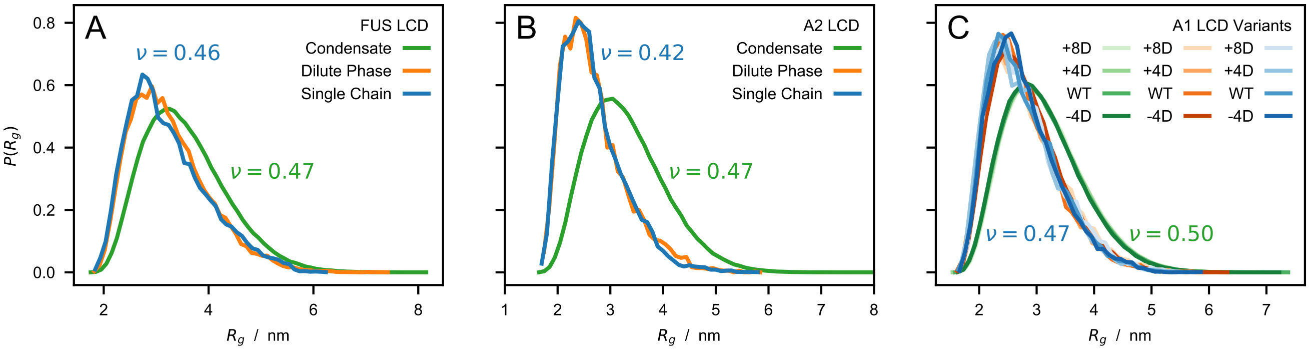


**Fig. S23.** Probability distributions of the radius of gyration, $R_g$, in condensed (green) and dilute (orange and blue) phases at 50°C of (*A*) FUS LCD, (*B*) A2 LCD, (*C*) wild-type and charge variants of A1 LCD simulated using the M1 model. Green and orange lines are calculated from coexisting protein-rich and dilute phases in multi-chain simulations whereas blue lines are calculated from single-chain simulations. Scaling exponents, $\nu$, are calculated for the single chain (blue) by fitting $R_{ij} = R_0|i-j|^\nu$ in the long-distance region, $|i-j| > 10$. For each protein, the best-fit $R_0$ values for the single chain are used to estimate $\nu$ from the average $R_{ij}$ of the chains in the condensate (green), with $\nu$ as the only free-fitting parameter.

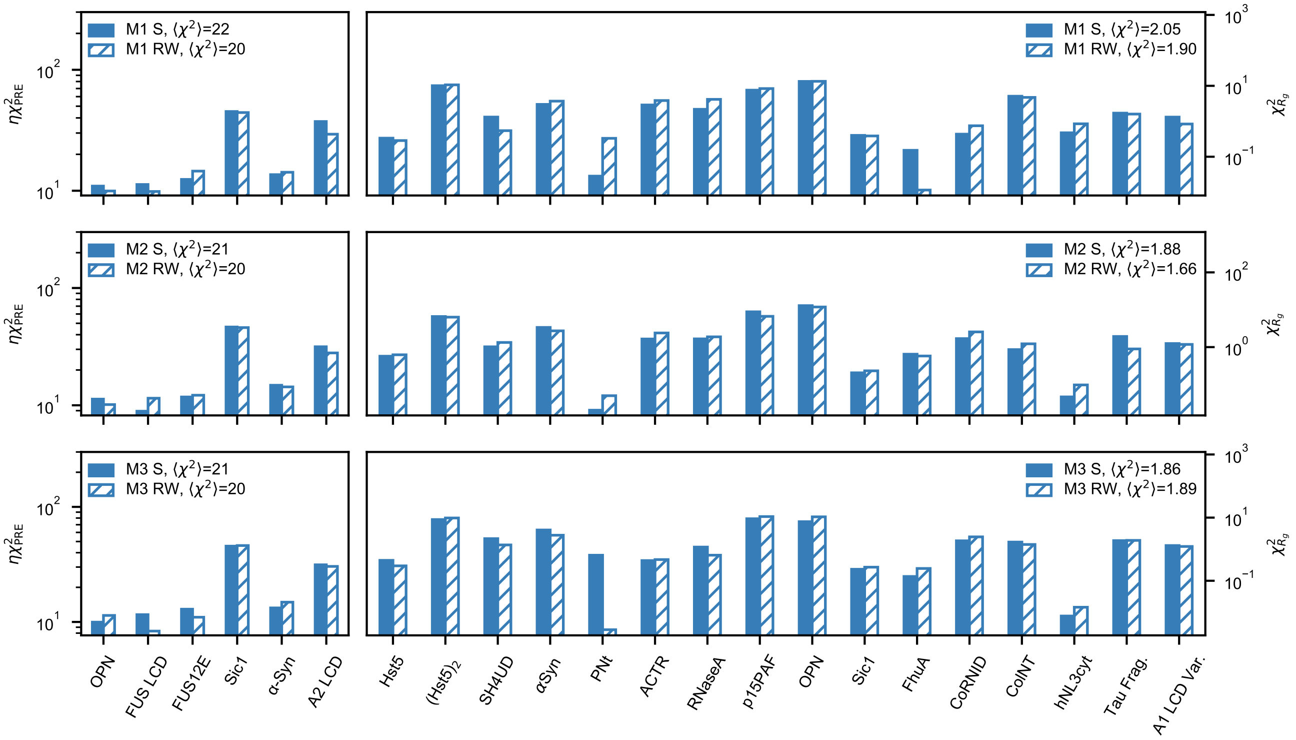


**Fig. S24.** Comparison of $\eta\chi^2_{PRE}$ and $\chi^2_{R_g}$ for the optimized models M1, M2 and M3, calculated directly from simulation trajectories (closed) and estimated by reweighting during the parameter-learning procedures (hatched).
